# Sequence and structural determinants of efficacious *de novo* chimeric antigen receptors

**DOI:** 10.64898/2025.12.12.694033

**Authors:** Arthur Chow, Hoyin Chu, Ruofan Li, Benan N. Nalbant, Abdul Vehab Dozic, Laura C. Kida, Caleb A. Lareau

**Author notes:** These authors contributed equally (Arthur Chow, Hoyin Chu).

## Abstract

Advances in generative protein design using artificial intelligence (AI) have enabled the rapid development of binders against heterogeneous targets, including tumor-associated antigens. Despite extensive biochemical characterization, these novel protein binders have had limited evaluation as agents in candidate therapeutics, including chimeric antigen receptor (CAR) T cells. Here, we synthesize generative protein design workflows to screen 1,589 novel protein binders targeting BCMA, CD19, and CD22 for efficacy in scalable protein-binding and T cell assays. We identify three main challenges that hinder the utility of *de novo* protein binders as CARs, including tonic signaling, occluded epitope engagement, and off-target activity. We develop computational and experimental heuristics to overcome these limitations, including screens of sequence variants for individual parental structures, that restore on-target CAR activation while mitigating liabilities. Together, our framework accelerates the development of AI-designed proteins for future preclinical therapeutic screening, helping enable a new generation of cellular therapies.

## INTRODUCTION

Chimeric antigen receptors (CAR) are synthetic proteins that redirect the specificity of immune cells toward a restricted set of antigens^1^, enabling the precision targeting of pathogenic cells that underlie cancer^2^ and autoimmunity³. Autologous CAR T cells have emerged as a transformative treatment modality for hematological malignancies, with seven FDA-approved therapies demonstrating robust clinical efficacy and durable patient responses^2–4^. The efficacy of tumor cell killing requires the design of synthetic binders that can sensitively and specifically recognize tumor-associated antigens (TAAs), and these binders are subsequently incorporated into CAR transgenes containing T cell activation domains^2^. Though most clinical-stage CARs have predominantly repurposed single-chain variable fragments (scFvs) for antigen recognition, in recent years, alternative protein binder formats have emerged and demonstrated promising preclinical efficacy by engaging TAAs and controlling malignant cell growth^5^.

Artificial intelligence (AI)-driven protein binder design is an emerging method for engaging TAAs, where novel protein binders are designed against target antigens *de novo*^6^. AI-designed proteins offer several theoretical advantages over traditional antibody-derived domains, including rapid generation *in silico*^7–9^, enhanced stability through optimized folding^10–12^, small size (∼60 amino acids), and prospective identification of the binding epitope^7,8,13^. These attributes of *de novo* protein binders stand to overcome efficacy limitations reported from various scFv-based CARs, including tonic signalling due to binder aggregation^14^ and enhanced immune synapse formation from rational epitope selection^15^. Motivated by recent work confirming the feasibility of AI-designed binders as antigen receptors in novel CAR formats^16–19^, we sought a comprehensive profiling of *de novo* designed proteins as efficacious antigen engagers, including an elucidation of the properties that elicit effective or ineffective CAR activation and signalling. Such a characterization could enhance the efficiency and developability of AI-designed proteins for cellular therapies and broaden the set of TAAs that could be targeted by CAR T cells.

Here, we evaluate the feasibility, structural correlates, and sequence determinants of efficacious CAR function with variable antigen-binding proteins created via generative protein design. We synthesize protein design pipelines that generate novel *de novo* binders against three antigens with clinical-stage scFv sequences as gold standards for benchmarking. Using a combination of scalable protein-binding and cell-based CAR assays, we study the binding and CAR signalling attributes of 1,589 distinct AI-designed TAA-binding proteins. Through this characterization, we identify three key constraints limiting *de novo* designed binders for CAR T cell efficacy, including tonic signaling, epitope occlusion from native ligands, and off-target binding. Critically, we demonstrate that these limitations can be overcome through rational protein design and engineering of non-interface residues. Our work provides a roadmap for developing next-generation CAR T cell therapeutics with enhanced efficacy and reduced liabilities, accelerating the utilization of *de novo* protein binders in future cell therapies across heterogeneous antigens.

## RESULTS

### De novo design of protein binders

We first assessed the feasibility of designing protein binders *de novo* against B-cell maturation antigen (BCMA), a surface protein overexpressed in multiple myeloma^20^ (**Fig. 1a**). We integrated various computational methods for two primary *de novo* binder design workflows. First, we implemented an RFdiffusion-based pipeline^9^, which spans backbone generation via RFdiffusion^8^, sequence diversification via ProteinMPNN^21^, and folding verification via AlphaFold2-multimer^22^. Second, we implemented BindCraft^7^, which generates protein binder sequences via protein hallucination by iteratively optimizing amino acid compositions using AlphaFold2 gradients. As part of our workflows, we applied the recommended tool-specific filters, such as interchain predicted alignment error (ipAE) and predicted local distance difference test (pLDDT), to ensure high confidence binding between the antigen target and designed *de novo* binders^9,23^ (**Methods**). For RFdiffusion, 20,000-30,000 structures were diffused targeting a known structure of the BCMA extracellular domain (Protein Data Bank, PDB: 1XU2). For BindCraft, we used the 4-stage multimer hallucination trajectory settings to design 1,000-2,000 binders satisfying default filtering criteria, which include other widely-adopted AlphaFold interaction metrics (e.g., Interface predicted template modelling; ipTM) as well as biophysical metrics, including the difference in solvent accessible surface area (dSASA; **Methods**). Once candidate molecules were generated and prioritized from these *in silico* metrics for both methods, synthetic DNA encoding 100-125 designed proteins (size 35-80 amino acids) was transformed as a pool into *S. cerevisiae* with a pETcon(-) yeast surface display (YSD) plasmid and individually into *E. coli with a* recombinant *E. coli* protein expression plasmid (**Extended Data Fig. 1a,b**). Binding was assessed in a pooled setting via flow cytometry of fluorescently labeled recombinant BCMA protein and quantitatively confirmed individually with recombinant protein via biolayer interferometry (BLI; **Fig. 1a; Methods**).

**Figure 1.**
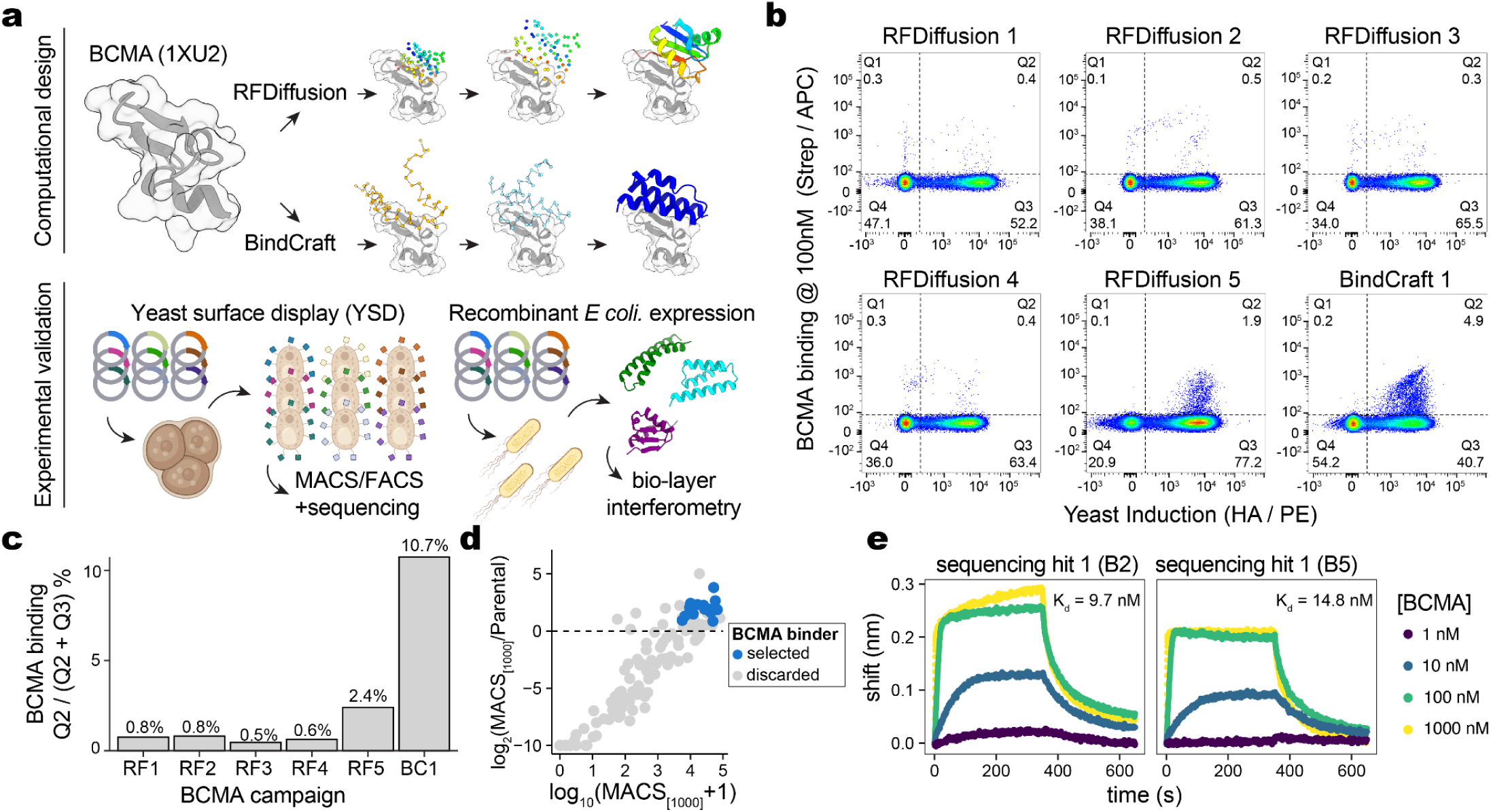
Summary of protein binder development using generative artificial intelligence. **(a)** Schematic of computational design (top) and experimental validation (bottom) methods for developing *de novo* binders targeting BCMA. **(b)** Yeast surface display (YSD) results of six experimental campaigns of binders for BCMA. **(c)** Quantification of YSD enrichment from (b). **(d)** Enrichment of individual binders from the BindCraft 1 (BC1) campaign, highlighting enriched binders selected for further validation. **(e)** Confirmation of binding for selected binders via biolayer interferometry (BLI).

To identify candidate BCMA binders, we executed six different design campaigns to target BCMA with YSD validation (**Extended Data Table 1; Methods**). For all yeast pools, we incubated yeast displaying protein minibinders with 100 nM BCMA and off-target antigens as control to verify the specificity of BCMA binding (**Fig. 1b; Extended Data Fig. 1c,d; Extended Data Table 2**). Across our six campaigns, we observed the largest enrichment with the singular BindCraft campaign relative to any of the five RFdiffusion campaigns, as 10.7% of induced yeast bound BCMA at 100 nM, compared to a maximum of 2.4% from the RFdiffusion campaigns (**Fig. 1c**). Magnetic-activated cell sorting (MACS) with recombinant BCMA protein was used to enrich yeast cells that bound the target antigen at variable concentrations before PCR-amplifying and sequencing the binder-encoding inserts (**Methods**). From our BindCraft campaign, we identified 17 sequences that showed compelling increase in abundance from MACS-enrichment and sequencing (**Fig. 1d; Extended Data Fig. 1e; Extended Data Table 3**). Recombinant *E. coli* expression and purification of 6 candidate sequences confirmed BCMA binding with affinities ranging from ∼2-200 nM (**Fig. 1e; Extended Data Fig. 1f**). While RFdiffusion produced a lower overall success rate of experimental binding, parallel workflows of YSD enrichment, sequencing, and individual sequence characterization via BLI confirmed BCMA binding for two candidate sequences from the RFdiffusion 5 campaign (**Extended Data Fig. 1g,h**). In total, our results support that both RFdiffusion and BindCraft can feasibly generate zero-shot *de novo* protein binders with affinities comparable to chimeric antigen receptors used in clinically validated CAR therapies. We conclude that BindCraft provides a novice-friendly workflow for rapid and facile generation of *de novo* protein binders, consistent with reports from public binder design competitions^7,23^.

### Characterization of CAR activities

Following the characterization of our *de novo* anti-BCMA binders, we sought to evaluate their performance as CARs using both CAR Jurkats as an *in vitro* cell model and primary human T cells for *ex vivo* assessment (**Fig. 2a**). We used a second generation CAR backbone with a 4-1BB co-stimulatory domain and fluorescent reporter protein (GFP) that enabled modular integration of new binders into the CAR backbone for testing (**Extended Data Fig. 2b; Methods**). We assessed tonic, off-, and on-target signaling by co-culturing CAR T and Jurkat cells with various tumor cell lines expressing variable, if any, levels of the BCMA target antigen (**Fig. 2a**). We selected CD69 surface expression on CAR^+^ cells as a primary readout, as CD69 is rapidly expressed following T cell activation, including CAR engagement^24,25^. We quantified the percentage of CD69^+^ cells among CAR^+^ cells, which enables a rapid and scalable assay for measuring CAR activity in Jurkats^25^ (**Extended Data Fig. 2a**). We compared the efficacy of our *de novo* binders to the C11D5.3 scFv, the anti-BCMA binder used in the FDA-approved CAR Abecma^26^.

**Figure 2.**
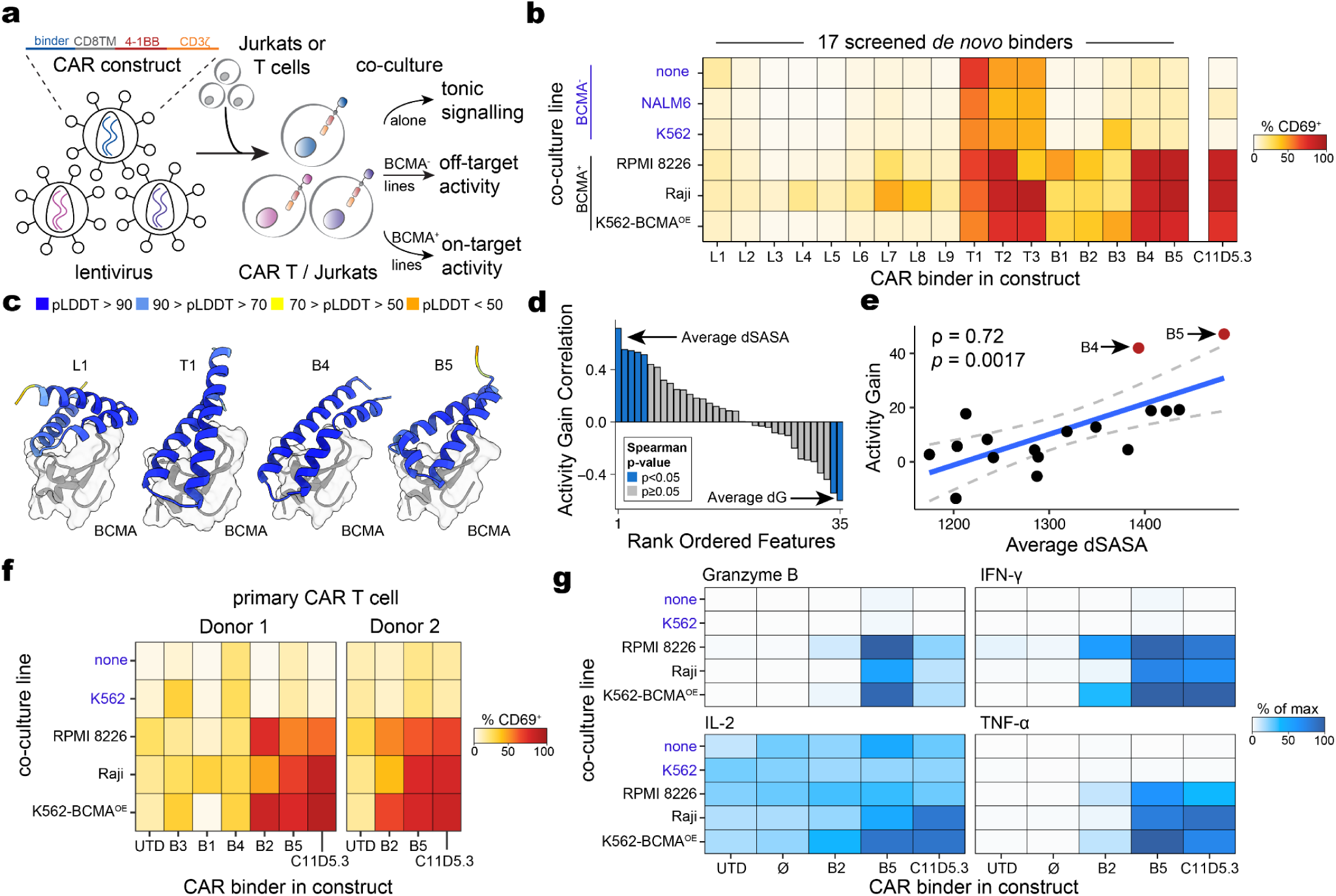
Characterization of *de novo* designed protein binders as CAR binders. **(a)** Schematic of CAR backbone and lentiviral-based systems for assessing CAR activity. **(b)** Screen of 17 *de novo* BCMA binders in heterogeneous conditions compared to the clinical scFv (C11D5.3). Shown is the %CD69^+^ Jurkats among GFP^+^ cells. **(c)** Predicted structures of four exemplar *de novo* CAR binders. Binders are colored by per-residue AlphaFold2 predicted Local Distance Difference Test (pLDDT) metrics. **(d)** Per-binder metrics associated with variable CAR function, highlighting the top two associated with CAR efficacy. **(e)** Summary of CAR activation efficacy against per-binder dSASA. **(f)** Activation profiles of *de novo* binders in primary CAR T cells with variable co-culture lines for two donors. **(g)** Cytokine production from *de novo* designed binders in primary CAR T cells (Donor 1).

After 16 hours of co-culture, we characterized the 17 binders from our BindCraft1 campaign, observing that 9 had low CAR CD69 activation (L1-L9; 2-43% CD69^+^ CAR^+^ cells), three featured high tonic signalling (T1-T3; 46-70%), and five had low tonic but high BCMA-dependent activation (B1-B5; 22%-90%) that were qualitatively similar to the gold-standard C11D5.3 scFv (72-85%; **Fig. 2b**). Notably, none of the binders showed meaningful evidence of off-target activity, which was assessed by comparing CD69 activation of BCMA-lines against the CAR Jurkats alone. All six designs were predicted to form as well-folded monomers with high-confidence *in-silico* quality metrics, and each predicted to fold into α-helical bundles with unique arrangements in complex with BCMA (**Fig. 2c**). For ease of interpretation, we defined activity gain as the difference in CD69 positivity between antigen-positive co-cultures and CAR-modified cells alone. This metric showed strong correlations with multiple *in silico* design metrics, most notably average dSASA (Spearman rho: 0.72, *p* = 0.0017; **Fig. 2d)**, with the top-performing binders B4 (79-85%) and B5 (77-90%) showing the highest overall BCMA-dependent CD69 activation (**Fig. 2e**). Moreover, we confirmed CAR activity from our two RFdiffusion-derived binders in analogous CAR Jurkat activation, consistent with recent reports^17,19^ (**Extended Data Fig. 2c,d**).

Encouraged by these results, we screened the five best Bindcraft binders (B1-B5) in primary CAR T cells **(Methods)**. We again observed strong BCMA-dependent CAR T activation, especially for the B2 (44-82% CD69^+^) and B5 binders (52-74% CD69^+^; **Fig. 2f)**. We replicated the clear antigen-activation result in a second donor, demonstrating overall comparable levels of primary T cell activation as the C11D5.3 scFv (66-82% CD69^+^; **Fig. 2f**). Next, we further assessed our two best *de novo* binders (B2 and B5) for hallmarks of productive CAR signalling response, including production of effector molecules and cytokines IL-2, IFN-γ, TNF-α, and Granzyme B following co-culture with the same cell lines via Enzyme-Linked Immunosorbent Assays (ELISAs; **Methods**). For all four molecules, binding of the B5 minibinder induced comparable or higher levels of production of effector molecules (IL-2: 29-100%; IFN-γ: 71-99%; TNF-α: 63-100%; Granzyme B: 52-100%) compared to the clinical-stage BCMA scFv (IL-2: 21-87%; IFN-γ: 64-98%; TNF-α: 38-88%; Granzyme B: 16-21%; **Fig. 2g**). Though our breadth of screening across variable constructs and conditions was predominantly performed as individual replicates, we verified consistency of key co-coculture readouts comparing B5 and the C11D5.3 scFv in triplicate (**Extended Data Fig. 2e**). In total, our characterization of antigen-dependent T cell activation and effector response confirms the feasibility of *de novo* minibinders as antigen receptors in CAR T cells, consistent with other reports of *de novo* binders targeting human and viral antigens^16–19^.

### Sequence variation underlies tonic signaling

In general, *de novo* designed minibinders that pass our chosen designability metrics fold into well-defined structures as predicted by state-of-the-art folding algorithms with high *in silico* confidence^9^. Unlike scFvs that engage antigens from complementary determining region (CDR) loops, *de novo* binders fold into rigid, stable structures that typically bind target antigens along non-polar residue sites. From these high-confidence backbones, algorithms like ProteinMPNN can generate hundreds of high-quality candidate sequences for an individual structure^21,27,28^. Hence, we reasoned that our design pipeline could sample sequence variation from a selected binder (i.e., B5), enabling correlative analyses to identify sequence and biophysical parameters that underlie efficacious CAR function.

To ensure the mutagenized binder sequences have high sequence variation while preserving high likelihood of BCMA binding and structural similarity, we developed the CAR-PNN (Chimeric Antigen Receptor-Protein-mpNN) workflow which diversifies the binder sequence at interface and non-interface residues and filters candidates based on BindCraft-inspired, Boltz1^29^-based filters. In particular, starting with our parental B5 binder, we applied ProteinMPNN trained with only soluble structures (SolubleMPNN)^30^ to generate a total of 4,000 sequences that mutagenized residues critical to the CAR synapse (<4Å) or that were not part of the binding interface (**Fig. 3a; Methods**).These candidate sequences were then refolded against BCMA via Boltz1^29^ and the average metrics across 5 diffusion samples such as pLDDT and dSASA were used for design filtering (**Methods**). Among the filtered sequences, we selected 21 additional CAR sequences that diversified over the number of interface hydrogen bonds, hydrophobic residues, charge, and dSASA with a minimum amino acid Levenshtein distance of six and maximum backbone RMSD of 1Å to the B5 parental (**Fig. 3b; Extended Data Table 3**). Overall, our selected binders had a median of 9 residues changed at the interface and median 21 changed from non-interface residues, with a total of 81% of possible residues mutated at least once (**Extended Data Fig. 3a**). Among our selected binders for, each had similar, if not improved, *in silico* performance metrics as the parental B5 binder, including complex interaction metrics correlated with performance of *de novo* binders^9^ (**Extended Data Fig. 3b**).

**Figure 3.**
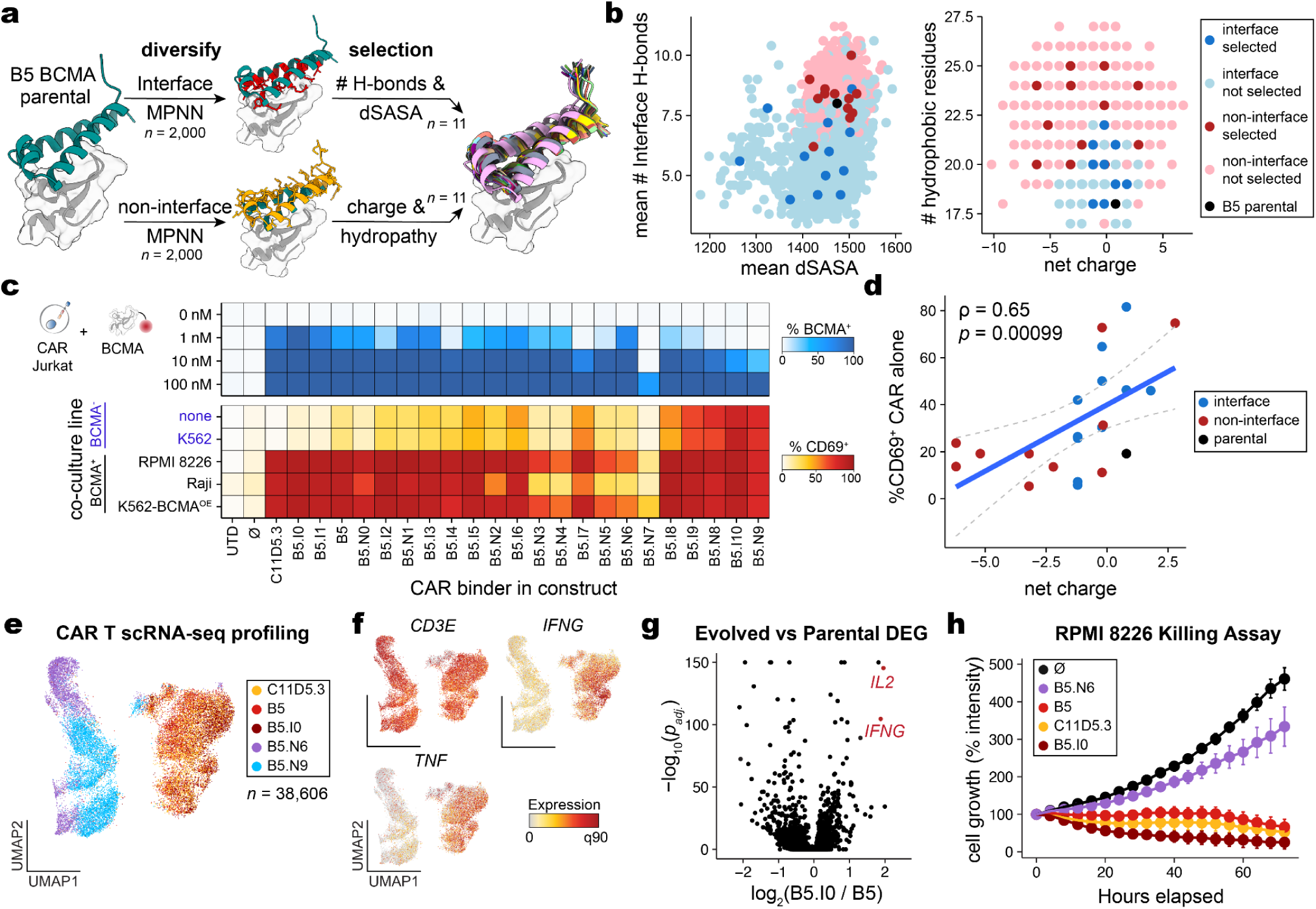
Sequence variation underlies CAR performance. **(a)** Schematic of diversifying BCMA binder sequences given a single binder (B5). **(b)** Summary of biophysical characteristics of diversified protein sequences, highlighting binders selected for additional CAR testing. **(c)** Summary of diversified sequences from antigen CAR flow (top) and co-cultures with variable cell lines (bottom). **(d)** Association of binder charge with tonic signalling (data from (c). *P value* and point estimate are reported from a Spearman correlation test. **(e)** UMAP embedding of 5 CARs profiled via scRNA-seq. **(f)** Marker gene expression across CAR T cells. **(g)** Summary of differentially expressed genes comparing evolved (B5.I0) and parental (B5) CARs. **(h)** Killing assays for selected CAR constructs against RPMI-8226, a BCMA^+^ line.

Synthetic DNA encoding each of these 21 binders were cloned into the same CAR construct, and CAR Jurkats were screened in parallel. First, to verify antigen binding, we added variable concentrations of fluorescently labeled recombinant BCMA protein to the CAR Jurkats and measured the percentage of CAR^+^ cells that bound the antigen (**Fig. 3c**). We observed binding of recombinant BCMA from all diversified sequences with modest variation at lower antigen concentrations. Co-culture of CAR Jurkats with BCMA-positive or negative cell lines showed considerable variance in CAR activation across conditions, including the B5.I0 sequence with improved activity relative to the B5 parental and comparable activity to the C11D5.3 scFv used in Abecma **(Fig. 3c; Extended Data Fig. 3c)**.

We observed 5 sequences with substantial tonic signalling and an overall activation gradient in this experiment ranging from ∼0 to ∼80% CD69 expression in the CAR-alone condition. As all proteins having similar predicted folds, confidence metrics, and BCMA binding, we attributed the variation in tonic signalling to underlying sequence properties of the BCMA binders. To assess the impact of the diversified biophysical and sequence metrics, we performed correlations of the tonic signalling and sequence attributes of these 22 CARs, observing a strong association between the binder net charge and the degree of tonic signalling (*r =* 0.65; *p =* 0.00099; **Fig. 3d**). Of note, no other biophysical parameter showed a strong correlation with CAR tonic signaling or activity gain, including hydrophobic content, number of hydrogen bonds, or dSASA **(Extended Data Fig. 3d)**. These analyses implicate binder net charge as a determinant of tonic signalling in *de novo* designed CAR T cells and a promising filtering heuristic for ensuring efficacious CARs. Our findings corroborate the report of positively charged patches (PCP) on scFvs as a major determinant of tonic signalling across 10 distinct antigens and binders^31^.

We next sought a more comprehensive characterization of our lead *de novo* binders that showed comparable T cell activation. To assess this, we first performed single-cell RNA sequencing (scRNA-seq) on primary CAR T cells expressing C11D5.3, the parental B5 binder, and three evolved minibinders (B5.I0, B5.N6, B5.N9) following co-culture with a BCMA^+^ cell line. The transcriptional profiles of the CAR T cells recapitulate the activation status seen in Jurkat screening, where cells expressing binders with strong antigen-dependent activation (C11D5.3, B5, B5.I0) clustered separately from those with minimal activity gain (B5.N6, B5.N9; **Fig. 3e,f; Extended Data Fig. 3e)**. Module scoring of CAR-T cells revealed strong effector activation accompanied by low exhaustion signatures for the C11D5.3, B5, and B5.I0 constructs, consistent with their high CD69 responses on BCMA^+^ target cells and low activation on BCMA^-^ cells **(Extended Data Fig. 3f)**. As expected, the cluster containing C11D5.3, B5 and B5.I0 overexpress effector cytokines such as *IFNG* and *TNF* **(Fig. 3f)**. In addition, cells with evolved binder B5.I0 had significantly higher expression of activation markers such as *IL2* and *IFNG* compared to the parental binder B5 **(Fig. 3g; Extended Data Fig. 3g; Extended Data Table 4)**. From these transcriptomic and cytokine-based cellular profiles, we hypothesized that our *de novo* CAR T cells would enable efficacious tumor cell control. Our screen of primary CAR T cells incorporating these different binders in *in vitro* killing assays demonstrated clear tumor control across a range of BCMA cell lines (-88.4% for CD11D5.3; 094.5% for B5.I0; **Fig. 3h)**. Compared to the parental B5 CAR and the scFv-based CAR, our evolved B5.I0 CAR further demonstrated enhanced tumor cell control across additional BCMA^+^ cell lines, including K562 overexpressing BCMA and Raji cells (**Extended Data Fig. 3h)**. Taken together, our analyses verified that selected AI-generated binders can activate T cell killing programs and control tumor growth with comparable efficacy for different sequences predicted to fold into near-identical binder structures.

### Variable antigen engagement in mammalian cells

A conceptually attractive aspect of computational binder design is the pre-specification of residues for binding on the target surface. For smaller targets like BCMA (36 amino acid extracellular domain), there are few potential contact residues, limiting the diversity of epitope binding. Conversely, for CD19, a B cell-restricted surface antigen targeted by multiple FDA-approved CAR T products for B cell leukemia and lymphoma, a much larger extracellular domain (∼280aa) results in many more candidate binding epitopes. For developing CAR T cells targeting CD19, prior work has indicated that binding epitopes in proximity to the tumor cell surface may be advantageous^32^. Hence, we evaluated the feasibility of designing binders against CD19, with a particular interest in targeting membrane-proximal epitopes that could enable more efficacious CAR activity.

To chart the binding geometry of existing clinical binders on CD19 relative to its endogenous co-factors, we superimposed the crystal structure of the widely used FMC63 scFv used in clinical CAR T cells in complex with CD19 (PDB: 7URV) to another crystal structure of CD19 in complex with its natural binding partner CD81 (PDB: 7JIC; **Fig. 4a**). Using these binding interfaces as a reference, we launched six distinct binder campaigns with variable hotspot residues on CD19 to develop minibinders (**Extended Data Table 1; Methods**). To our surprise, regardless of the input parameters, nearly all binders passing *in silico* design filters were constrained to the same CD81 binding epitope, likely reflecting that generative methods are prone to copying known protein complex interfaces and are biased toward hydrophobic patches (**Fig. 4b**). Despite the limited diversity of these interfaces, we performed YSD screening of synthesized binders, again observing an enrichment for CD19 binders developed via BindCraft (BC2, 9.9%) with no clear evidence of binding for any of the RFdiffusion campaigns (RFD1-4, 1-3%; **Fig. 4c-d**). The major difference between campaigns BC1 and BC2 was the overall size of the binder, which we scaled from 50-65 amino acids to 100-200 amino acids (**Methods**). While noting that emerging protein design methods may enable targeting geometric matches^33^ and disordered residues^34,35^, we observe that the current generation of widely-used binder design tools are limited in their ability to position binders onto physiologically appropriate surfaces for some target antigens.

**Figure 4.**
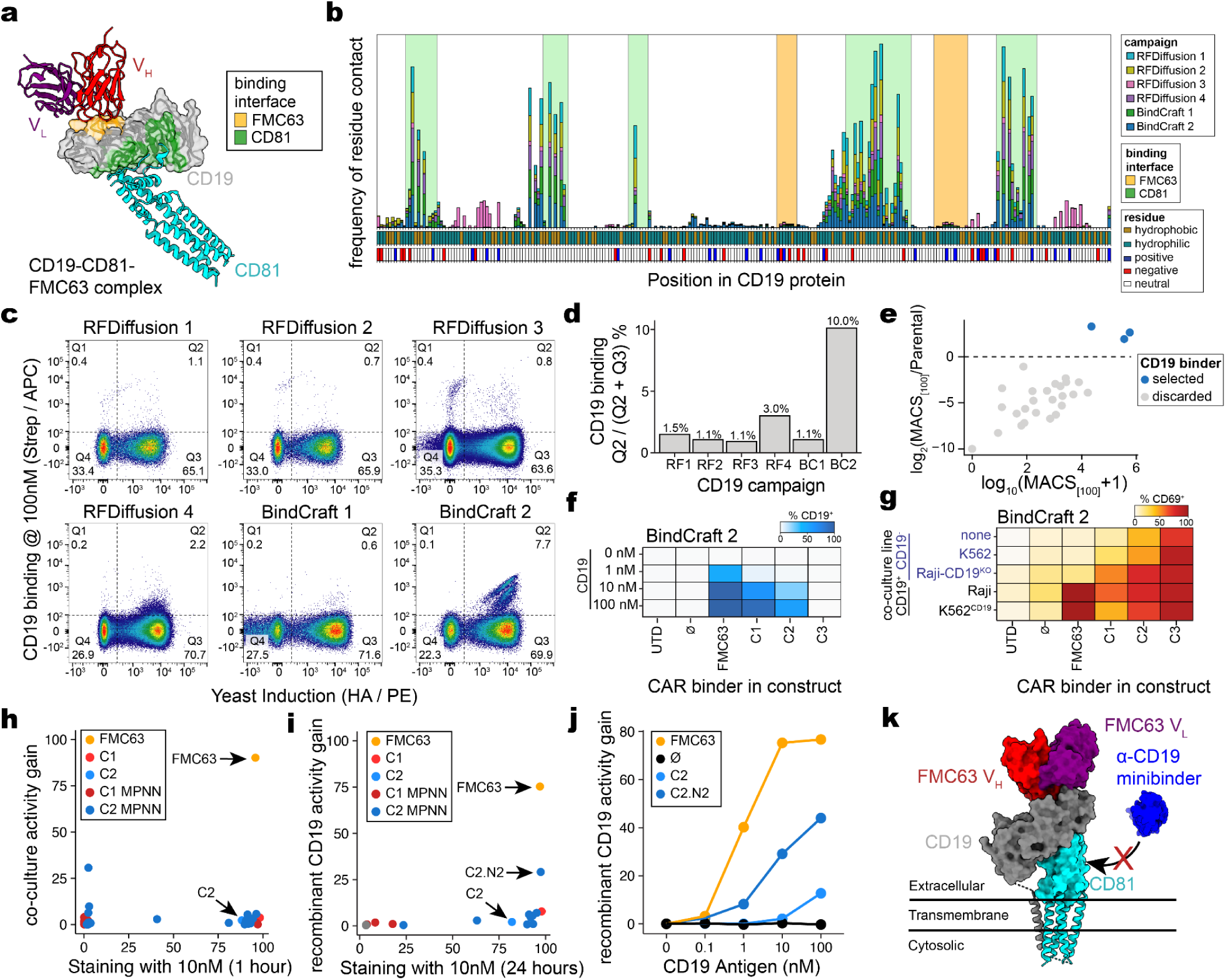
Native binding partners occlude binding epitopes from *de novo* CD19-directed CARs. **(a)** Structural rendering of the FMC63-CD19 complex (PDB: 7URV) aligned to the CD19-CD81 complex (PDB ID: 7JIC). The binding surfaces on CD19 for both proteins are highlighted. **(b)** Summary of CD19 protein, highlighting residues and binding surfaces. Barplots show the normalized proportion (out of 100 draws) of contacted residues from design campaigns. **(c)** Summary of yeast surface display binding of *de novo* designed proteins against CD19. **(d)** Quantification of YSD enrichment across six campaigns. **(e)** Quantification of individual binders enriched from the BindCraft 2 (BC2) campaign, highlighting enriched binders selected for further validation. **(f)** Characterization of CAR antigen binding at variable recombinant CD19 concentrations. **(g)** Overview of CAR Jurkat activation screening from BindCraft 2 campaign. **(h)** Summary of CD69 activation via coculture of C1 and C2 CARPNN evolved binders with Ramos cell line, highlighting selected CARs. **(i)** Same as (h) but for CAR activation via 24 hours of recombinant CD19 incubation. **(j)** Concentration-specific activation of variable CARs with recombinant CD19 antigen. **(k)** Schematic of occluded epitope engagement of CD19-directed minibinders from CD81 binding on target cells using the aligned FMC63-CD19-CD81 complex.

From our BC2 campaign, we identified three enriched sequences that bound to CD19, which we subsequently expressed into CAR Jurkat cells (**Fig. 4e-g**). For one CD19 binder, C3, we did not observe recombinant CD19 binding when expressed as a CAR in Jurkat despite a strong pooled enrichment following MACS (**Fig. 4f**). Despite clear evidence of CD19 binding in YSD, our results suggest that this simplified system for protein binding may not translate into human cells, and thus we recommend staining CAR Jurkats with fluorescently-labeled recombinant antigen to discriminate binders subject to this limitation. Otherwise, co-cultures with the enriched binders C1 and C2 showed limited CAR activation despite their clear binding to recombinant antigen (**Fig. 4g**).

Motivated by our success diversifying the underlying sequences from our parental BCMA binder, we reasoned that our CARPNN workflow could produce sequence variants with improved CAR activity **(Extended Data Fig. 4a-f**). However, consistent with the limited CAR activity observed in our initial Jurkat co-culture screen, the CARPNN-evolved binder C1 and C2 variants did not show any measurable improvement. Notably, mutations within the binding interface fully ablated CD19 binding, suggesting that the CD81-associated epitope is geometrically constrained and affords minimal tolerance for sequence variation.

To evaluate functional engagement, we compared recombinant CD19 binding of each construct with its CD69 activation across CD19⁺ tumor cell lines. FMC63-expressing CAR cells displayed both high antigen affinity (96% binding at 10nM CD19) and potent CAR activation (90.2%, CD69 activity gain), whereas all evolved binders fell substantially below this benchmark, indicating a general decoupling between apparent CD19 binding and functional cellular activation (**Fig. 4h**). Further testing against free, soluble antigen alone revealed that one variant, C2.N2, induced clear antigen-dependent CD69 activation (29%, CD69 activity gain; **Fig. 4i,j**), strongly suggesting that the lack of function in cell-based assays is due to epitope occlusion by the native binding partner, CD81. Together, these data indicate that although several designed binders can recognize recombinant CD19, the dominant CD19 epitope targeted by the *de novo* binders overlaps with the region occluded by CD81 in the native CD19-CD81 complex (**Fig. 4k**), highlighting an additional challenge for therapeutic *de novo* design.

### Identifying and rescuing off-target CAR activity

Having established functional CAR binders targeting the two antigens with FDA-approved products, we sought to develop CAR-compatible minibinders against an earlier-stage target, CD22^36,37^. RFdiffusion and BindCraft campaigns **(Extended Data Table 1)** yielded protein binders with strong *in silico* success metrics against a hydrophobic patch within domain 7 of CD22 (**Fig. 5a**). YSD and sequencing again confirmed the high experimental success rate of BindCraft (1.9%), resulting in four candidate binders (D1-D4) that were screened using our CAR Jurkat workflow (**Fig. 5b,c**). Next, recombinant antigen flow and on-target CAR activation against CD22-expressing cells confirmed on-target activation with minimal tonic signalling (**Fig. 5d**). Surprisingly, we also observed a gain in CD69 expression (42.9% and 59.6%) in co-cultures with the RPMI-8226 line, a verified CD22^-^ cell line, for the D1 and D2 CARs, respectively (**Fig. 5d,e**). Conversely, RPMI-8226-dependent CD69 activation did not occur for the CAR expressing m971 (6.16%), the clinical-stage scFv targeting CD22.

**Figure 5.**
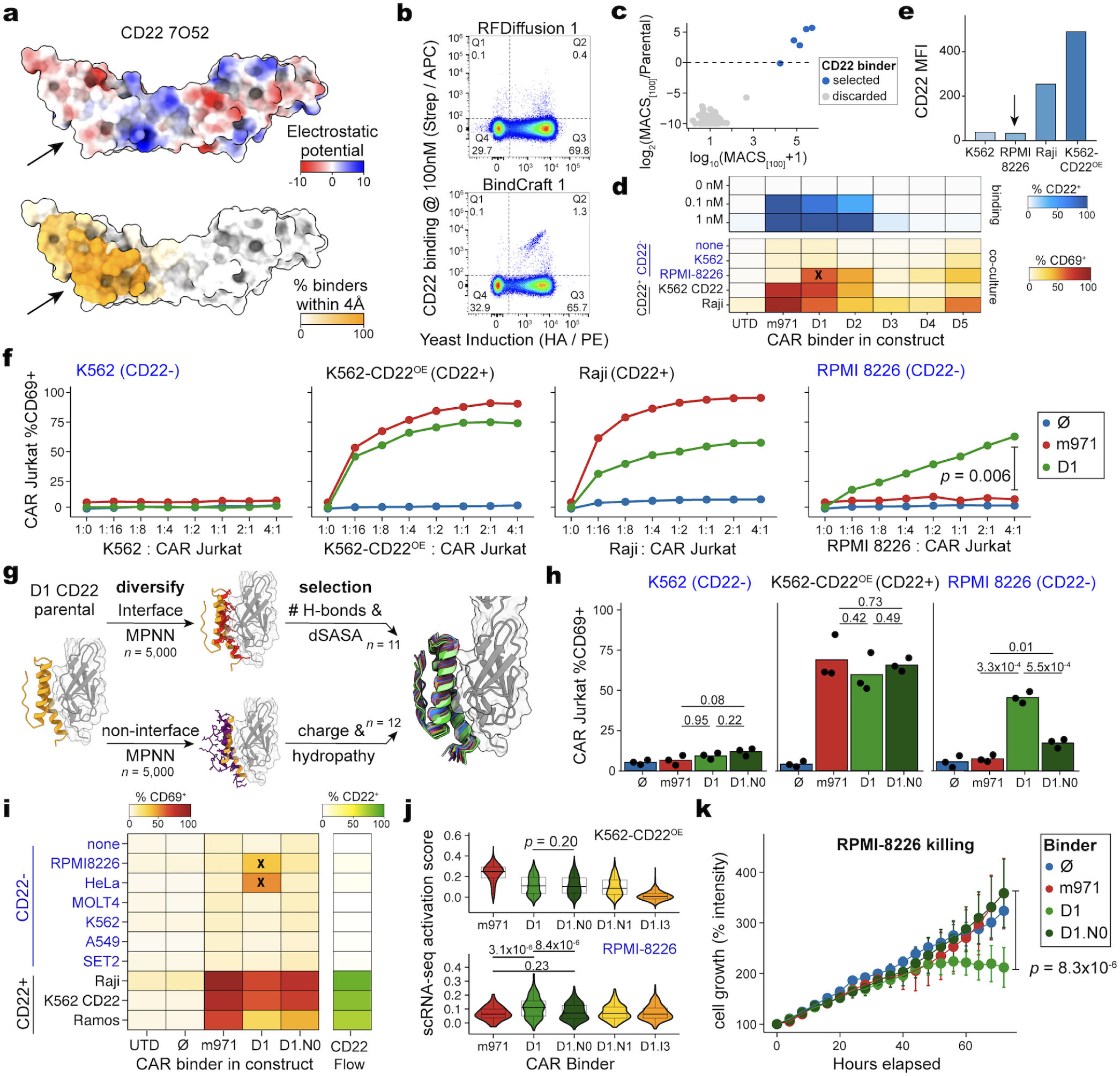
Reverting off-target binding through engineering *de novo* designed CD22-directed CARs. **(a)** Schematic of CD22 target antigen, with the arrow highlighting the preferred binding pocket as a hydrophobic patch. **(b)** Summary of YSD screening of *de novo* designed proteins from two campaigns against CD22. **(c)** Identification of four hits from the BindCraft campaign via sequencing. **(d)** CAR co-culture of four *de novo* CD22 binders (D1-D4) compared to m971 (clinical CAR). Shown is the %CD69^+^ Jurkats among GFP^+^ cells. **(e)** Validation of CD22 expression of cell lines in the CAR co-culture. Arrow highlights the absence of CD22 expression in RPMI 8226. **(f)** Cocultures of three CARs with variable effector to target (E:T) ratios. Statistical test: Wald test of linear regression comparing D1 *de novo* binder to m971 clinical CAR, adjusting for E:T ratio. **(g)** Diversifying CD22 binder sequences given a single binder (D1). **(h)** Triplicate CAR Jurkat co-cultures with variable CAR binders. Statistical test: Two-sided Student’s *t* test. **(i)** Summary of diversified CD22 sequences in CAR co-culture. “X” highlights off-target activation from parental binder, D1. **(j)** Activation scores from scRNA-seq profiles of five CAR binders cultured against two different cell lines. Statistical test: two-sided Mann-Whitney U test. **(k)** Primary CAR T killing curves against RPMI 8226 (CD22^-^) showing off-target-specific killing in the *de novo* D1 binder. Statistical test: Wald test of linear regression interaction term between D1.N0 binder and time compared to D1, adjusting for time and binder.

To verify the putative off-target activation from the D1 binder in the RPMI-8226 co-cultures, we repeated the CAR Jurkat co-cultures with variable effector-to-tumor (E:T) ratios (**Methods**). The variable co-culture compositions yielded the expected results for CD22^-^ and CD22^+^ lines, where Jurkat activation progressively increased overall when in culture with greater abundance of CD22^+^ cells (**Fig. 5f**). For RPMI-8226, we similarly observed dose-dependent activation of the D1 *de novo* minibinder (up to 65%) but not the m971 scFv (max 11%; *p* = 0.006; two-sided Wald test from linear model), confirming off-target activation from co-culture with a CD22^-^ line.

We hypothesized that our CARPNN workflow could reduce this off-target liability while retaining on-target activity. To achieve this, we generated and scored 10,000 diversified sequences based on the D1 binder, selecting an additional 23 binders for screening in CAR Jurkats for on- and off-target activation (**Fig. 5g; Extended Data Fig. 5a**). Remarkably, evolved sequences with sequence variation outside of the CD22 binding interface resulted in more specific CD22-dependent CAR signaling (**Extended Data Fig. 5b,c**). In particular, we observed one variant, D1.N0, that retained strong on-target activity, exhibiting strong recombinant binding (78% of CAR cells bound at 0.1 nM CD22), low tonic signaling (18% tonic CD69), and CD22-specific activation (65-74% CD69 activation) comparable to both m971 (83% CAR cells bound at 0.1 nM CD22; 5% tonic CD69; 61-90% CD22-specific CD69) and the parental D1 binder (64% CAR cells bound at 0.1 nM; 11% tonic CD69; and 55-61% CD22-specific CD69) However, the non-interface mutagenesis, eliminated off-target activation from co-cultures with RPMI-8226 (D1: 42% vs D1.N0: 19%), which was verified in triplicate (**Fig. 5h; Extended Data Fig. 5f**). To further verify this result, we expanded our co-culture screening across additional CD22^-^ cell lines to assess any unanticipated off-target activity. Among these, co-culture with HeLa cells again showed evidence of CD22-independent CD69 activation (48.8%) for the parental D1 binder but not the clinical scFv (7.8%) or the evolved D1.N0 variant (8.2%) **(Fig. 5i)**. Moreover, scRNA-seq profiling of primary CAR T cells encoding various minibinders confirmed on-target activation of m971, D1, and two evolved variants (D1.N0 and D1.N1), but activation was restricted to the D1 parental binder when in co-culture with RPMI-8226– both observations consistent with the Jurkat data (**Fig. 5j; Methods**). Consistent with these activation signatures, the D1.N0 CAR-T preserved cytotoxicity against CD22^+^ targets (**Extended Data. 5c,e**), while substantially limiting off-target killing of CD22^-^ cells (**Fig. 5k**), accompanied by cytokine profiles that were appropriately antigen-dependent and minimized in CD22^-^settings (**Extended Data Fig. 5d**).

To resolve potential off-target interactions driving the parental D1 binder’s nonspecific activation, we intersected the Human Protein Atlas (HPA) surfaceome with GTEx expression data to nominate candidate antigens with expression patterns matching the observed off-target activation profile (**Methods**). By selecting genes highly expressed in RPMI-8226 and HeLa cells but minimally expressed in K562, SET-2, and A549 cells, we narrowed the full surface proteome to a focused set of 19 plausible off-target candidates (**Extended Data Fig. 5g)**. Among these candidates, the chemokine receptor CXCR4 emerged as a notable possibility, as the parental D1 binder exhibited a markedly higher ipSAE^38^ score (0.66) of *in silico* co-folding toward CXCR4 than the D1.N0 variant (0.08; **Extended Data Fig. 5h**), consistent with reduced off-target interaction predicted for this antigen. However, as the predicted D1 binding site resides on a cytosolic portion of CXCR4, it is unlikely that this domain is accessible to CAR T recognition (**Extended Data Fig. 5i**).

Together, our characterization both motivates the need for extensive off-target screening of *de novo* designed proteins and provides a facile strategy for mitigating off-target activation via sequence diversification and screening. Our observations highlight the inherent complexity of off-target prediction for *de novo* binders, where structural plausibility must be reconciled with the physiological accessibility of candidate epitopes. As the rescuing mutations were located at non-interface residues and there is a lack of credible *in silico* predicted interaction candidates, we also highlight the off-target interaction could occur in non-interfacial residues, a paradigm that, to the best of our knowledge, is rarely utilized in conventional antibody engineering but has been corroborated by antibody language models^39^.

### Inference of efficacious CAR designs from *in silico* attributes

We hypothesized that a retrospective analysis of our campaigns could aid in identifying *in silico* heuristics for prioritizing sequences with a higher success rate, both for protein binding and CAR activation. First, we wondered whether the consistent improved performance by BindCraft relative to RFdiffusion could be explained across our three antigen campaigns. Upon compiling the binder sequences, we identified a clear increase in alanine composition among binders generated by the RFdiffusion workflow compared to BindCraft **(Fig. 6a)**. We reason that the overrepresentation of alanine creates poorly packed hydrophobic interfaces with largely nonfunctional side chains, reducing stability and likely contributing to the weak or absent binding from binders created in the RFdiffusion campaigns despite passing other recommended *in silico* success metrics. Subsequently, using Boltz-1^29^, we computed all binder sequences that we have tested in YSD and/or CAR activation assays to determine relative optimal metrics that have emerged across design tasks, including ipTM^7^, pAE interaction^9^, and ipSAE^38^. To quantify the binding affinity of binders that have been tested in Jurkats, we estimated the Kd based on a non-linear least square (NLS) fit of the staining intensity at different concentrations scaled to the known affinity of the positive controls reported in other studies^40–42^ (**Methods, Extended Data Fig. 6a,b**). We found that for binders tested in the CARPNN campaigns, while decrease in ipAE and increase in ipTM generally corresponded to higher affinity binders, ipSAE was the best calibrated cutoff metric where binders with ipSAE higher than 0.85 had the highest success rate at achieving sub 1000 nM binding across different campaigns (**Figure 6b, Extended Data Fig. 6c**). Similar behavior was also seen in our yeast surface display campaigns, where a single ipSAE cutoff of 0.85 was consistently the best single cutoff metric across different campaigns (**Figure 6c, Extended Data Fig. 6d,e**). For example, 4.35% (*n* = 3/95) of the tested CD22 binders which all passed the default BindCraft filter showed enrichment at 1000nM by YSD. When retrospectively applying the additional ipSAE ≥ 0.85 filter, only 10 candidates would have passed the threshold, including 30.0% (*n* = 3/10) which showed enrichment at 1000nM by YSD. Thus, our analyses strongly support the inclusion of ipSAE as a filtering metric for *in silico* prioritization in parental and ProteinMPNN-based evolution campaigns.

**Figure 6.**
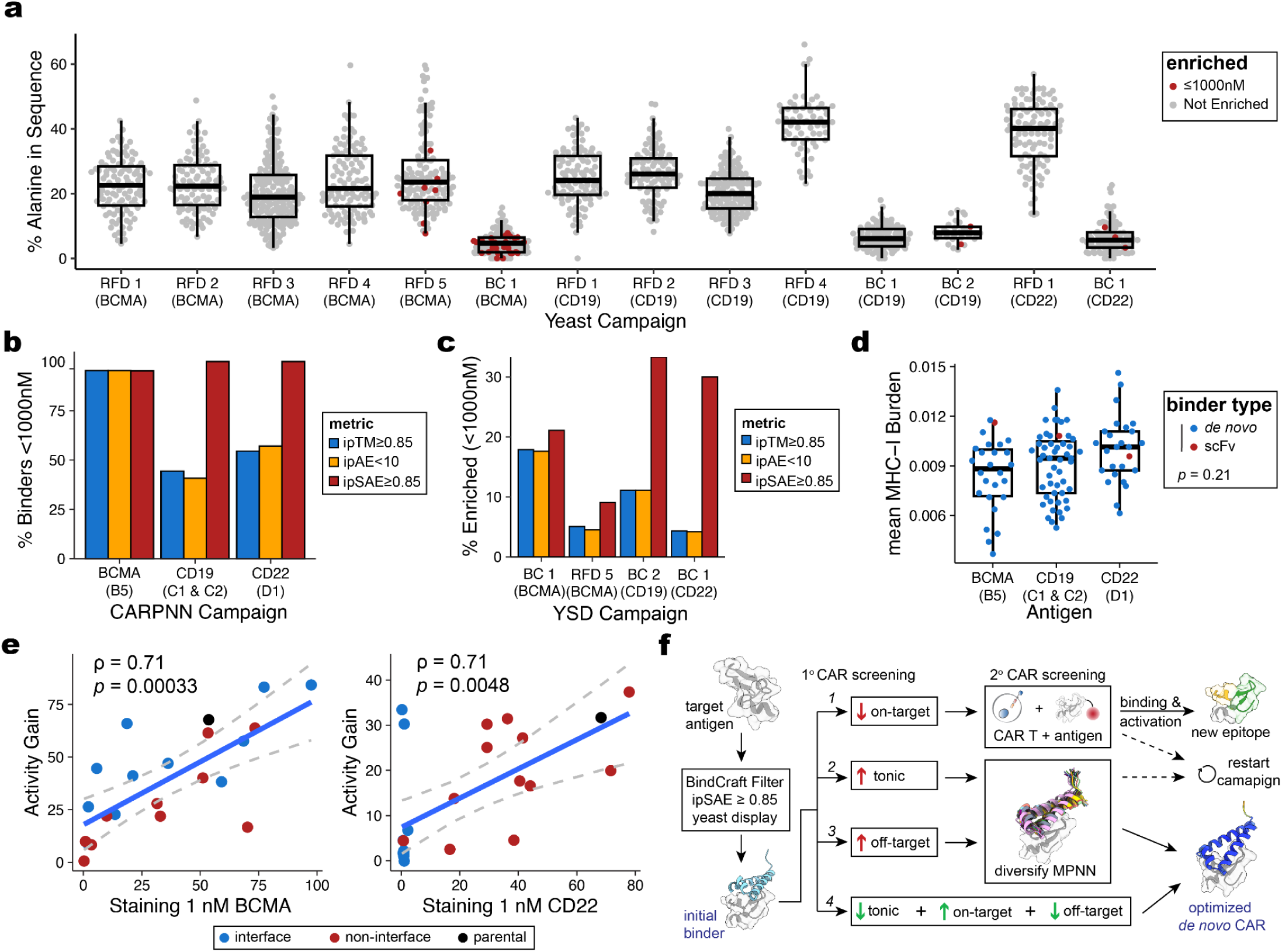
Retrospective campaign analysis reveals *in silico* correlates of successful CAR design. **(a)** Percentage of alanine in binder sequences tested in yeast display campaigns: (RFD, RFdiffusion; BC, BindCraft) **(b)** Proportion of binders within the CARPNN workflow with binding affinity stronger than 1000nM using various *in silico* cutoffs across three antigens. **(c)** Proportion of binders enriched in yeast surface display campaigns with binding affinity stronger than 1000nM using various *in silico* cutoffs. **(d)** Comparison of the average fraction of peptides predicted to bind strongly to class I MHC-I across all alleles supported by NetMHC 4.2^44^. Statistical test: Wald test statistic from generalized linear model. **(e)** Comparison of staining intensity via recombinant antigen at 1nM compared to mean proportion of CD69^+^ CAR+ cells cocultured with antigen-positive cells subtracted by the proportion CD69^+^ CAR^+^ cells alone. **(f)** Schematic of recommended workflow for designing *de novo* minibinder-based CAR.

Beyond binding metrics, we sought to understand potential sequence liabilities in our *de novo* binders that could limit long-term efficacy in more complex settings. As anti-CAR antibodies have been reported from multiple products that utilize scFvs^43^, we sought to assess the immunogenic potential of our *de novo* binders from the underlying binder sequences using NetMHC^44^ (**Methods**). We found that when normalized for length, *de novo* binders do not have higher likelihood of containing immunogenic peptides than existing CAR scFvs (**Fig. 6d; Extended Data Table 5**). Owing to the small size of our *de novo* protein binders, we further observed fewer predicted immunogenic peptides across diverse HLA alleles compared to corresponding clinical scFv (**Extended Data Fig. 6f**). Though *in vivo* testing is required to assess the immunogenicity of *de novo* protein binders compared to scFvs, we do not observe evidence of increased sequence liabilities. Further, as our work demonstrates considerable sequence diversity can underlie similar CAR structures with similar functional properties, we envision future design campaigns may apply additional sequence constraints to minimize immunogenicity while retaining efficacy.

Finally, an open question in developing protein binders for use in CARs lies in determining the optimal affinity window that maximizes functional activity. Clinical data from pediatric patients treated with a low-affinity CD19 CAR demonstrated that a 14 nM scFv outperformed the picomolar-affinity FMC63 scFv (∼0.3nM), suggesting that moderate affinities can yield superior T cell expansion, safety, and antitumor activity^45^. Using our successful binders from the BCMA and CD22 campaigns, we directly compared the staining intensity of the antigen at 1nM to CAR activity gain (**Methods**). We observed a strong correlation between the staining intensity and activity gain for both BCMA (*p*=0.00033; two-sided Spearman correlation test) and CD22 (*p*=0.0048; two-sided Spearman correlation test), suggesting that improved affinities of *de novo* binders generally enhances CAR efficacy for these proteins (**Fig. 6e**). In contrast to scFvs that can produce picomolar affinity binders from immunization campaigns, current tools that produce binders on the order of 1-1,000 nM likely benefit from affinity optimization. Combined with our successful application of ProteinMPNN-based redesigns once a parental structure binder has been identified, we suggest that sequence evolution, particularly at non-interface residues, can simultaneously improve affinity and CAR activity while mitigating liabilities, including immunogenic epitopes, off-target binding, and tonic signalling.

## DISCUSSION

Generative artificial intelligence has accelerated the adoption and success of *de novo* protein design, enabling the rapid creation of novel structures with potential therapeutic properties at a high success rate^6,46^. In particular, the success of diffusion and hallucination methods to generate binders against heterogeneous antigens reinforces *in silico* design as a compelling alternative to conventional modes of binder discovery, including antibody discovery campaigns from large library screens and/or immunizations. Hence, a straightforward application of *de novo* binders is the stand-in replacement of antibody fragments in these established modalities, including drug conjugates, bispecifics, and cell therapies. However, there have been limited reports guiding considerations of protein sequence design that would enable preclinical efficacy of designed proteins beyond recombinant antigen binding. Through our assessment of 1,589 binding proteins and engineering of CARPNN, our study outlines critical considerations for enhancing the efficiency of protein binder design and their use as CARs.

To generate parental protein binders *de novo*, we focused on the application of two leading binder design tools, RFdiffusion and Bindcraft. Across our campaigns of three different target antigens, we observed a consistently higher binder design success rate in campaigns where BindCraft was used as the primary design method. This can be largely attributed to the more stringent filtering criteria implemented by BindCraft, as we have shown in the CARPNN campaigns where a similar filtering criteria implemented in Boltz is able to also achieve a high binding success rate. In addition, BindCraft’s performance in community-based benchmarks^23^ and its ease of use allowed for rapid comparison and facilitated fast adoption, highlighting the importance of open benchmarks and tool accessibility in protein design. In the spirit of these principles, our study presents one of the largest datasets of *de novo* binders measured using a standardized YSD across multiple concentrations. In addition to the data presented in our study, we also propose using ipSAE as an *in silico* metric for binder identification. While other studies have also identified ipSAE as an effective metric for classifying binders from meta-analysis of other published *de novo* datasets, the definition of binding varies across studies and to our knowledge, this is the first time ipSAE is shown to be a calibrated metric across antigens with an explicit cutoff value, which could be informative for future design campaigns.

To enable AI-designed proteins for future candidate therapeutics, further characterization is required beyond recombinant protein binding. However, little guidance has been reported for efficacious binders in creating immune synapse for T cell-mediated killing. Through engineering our BindCraft-derived binders against our target antigens, we identified three limitations and recommendations to overcome them (**Fig. 6f**). First, through diversifying the parental BCMA binder sequence, we identify that higher numbers of positive residues are associated with increased tonic signalling, consistent with a report of positively charged residues in scFvs^31^. Unlike prior characterizations considering 10 different target antigens, our framework uniquely controls for similar structural, antigen-binding, and approximate antigen affinity to assess sequence-to-function relationships of CAR activity, including tonic signalling. Second, by characterizing CD19-directed CARs, we observed CAR-mediated activation from recombinant protein binding but not cell co-cultures, reflecting that our *de novo*-designed binders could not activate against cellular CD19. This discrepancy suggests that our designs targeted an epitope on CD19 that is masked by CD81 in its native heterodimeric complex, which underscores a fundamental limitation of the current generative design tools. Namely, *de novo* design methods may bias interfaces toward geometrically tractable surfaces rather than physiologically accessible epitopes on native multimolecular complexes. We anticipate that other interfaces may be occluded by features such as post-translational modifications (e.g., protein glycans), thereby inhibiting on-cell binding by *de novo* proteins. Finally, through extensive co-culture of our CD22 CARs, we characterized off-target activation from co-cultures with CD22^-^ cell lines. However, by extending our CARPNN framework, we could abrogate off-target signalling through sequence evolution of the non-interface binding residues. As co-folding analyses could not identify the putative off-target, we recommend extensive co-culture screening for off-target activation and subsequent sequence evolution to mitigate off-target activation in *de novo* CARs.

Looking forward, we expect that our characterization of challenges and limitations of *de novo* designed molecules will improve the efficiency of developing candidate molecules with improved efficacy and reduced liabilities. These efficacious binder design principles will further enable screening more complex T cell circuits, including multi-specific formats via AND^47^, IF-THEN^48^, and NOT^49^ CAR activation. We further envision an integrated loop where comprehensive single-cell profiles of primary diseased tissues nominate optimal combinations of antigens^50^ that can be, in turn, rapidly synthesized into sensitive and specific cellular therapies enabled by generative artificial intelligence.

## EXTENDED DATA TABLES

**Extended Data Table 1. Summary of yeast surface display campaigns and parameters.**

**Extended Data Table 2. List of antibodies for flow cytometry**

**Extended Data Table 3. Summary and annotation of all tested protein binders.**

**Extended Data Table 4. Differentially expressed genes from *de novo* CAR T cells.**

**Extended Data Table 5. NetMHC epitope predictions for CAR binders.**

## ACKNOWLEDGEMENTS

We are grateful to J. Gutierrez, E. Cumming, J. Chew, C. Bracken, W. Lin, and other members of the Lareau Lab for helpful feedback and discussions. A.C. is supported by a Basic and Translational Immunology Postdoctoral Award granted by the Ludwig Center at Memorial Sloan Kettering. This work was supported by NIH grants R00HG012579, R33CA302491, the MSKCC Cancer Center Grant P30CA008748, and an AML SPORE P50CA254838 award. C.A.L. is also supported by a National Academy of Medicine Catalyst award, the Michelson Prize for Immunology, and the Geoffrey Beene Cancer Research Center of Memorial Sloan Kettering. The funders had no role in study design, data collection and analysis, decision to publish, or preparation of the manuscript.

## AUTHOR CONTRIBUTIONS

A.C., H.C., and C.A.L. conceived and designed the study. A.C. led experimental design and execution with support from R.L., B.N., L.C.K., and A.D. H.C. led computational analyses with input from A.C. and C.A.L. A.C., H.C., and C.A.L. wrote the manuscript with input from all other authors.

## CODE AVAILABILITY

Custom code for replicating downstream analyses is available online https://github.com/clareaulab/denovo-cart-reproducibility. The method for diversifying and scoring CAR minibinder sequences is available on GitHub https://github.com/clareaulab/CARPNN.

## DATA AVAILABILITY

Raw and processed single-cell RNA sequencing of *de novo* and scFv CAR T cells is available at **GSE310399** with reviewer access code **ixapqscmnxwbdod**.

## FIGURES

**Extended Data Figure 1.**
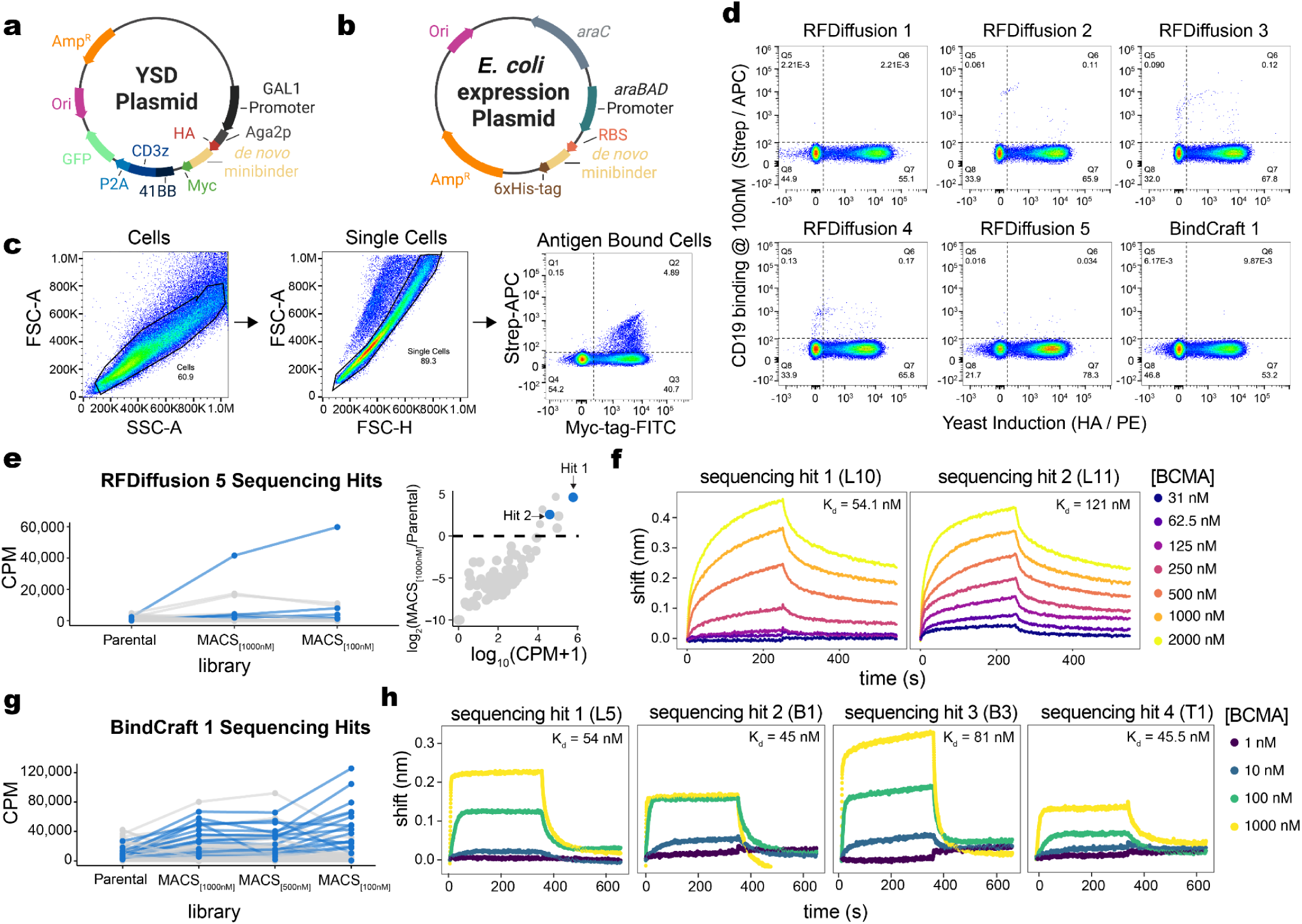
Characterization of lead *de novo* designed BCMA binders. **(a)** Yeast surface display plasmid encoding a *de novo* minibinder flanked by an N-terminal HA-tag and a C-terminal Myc-tag under a GAL1 promoter. **(b)** *E.coli* protein expression plasmid encoding *de novo* minibinder flanked by a 6xHis-tag for purification under a araBad promoter. **(c)** Representative gating strategy used during yeast display campaigns to identify binder hits. **(d)** Decoy binding of designed binder campaigns against CD19 antigen (100 nM), demonstrating variable levels of off-target reactivity. **(e)** Enrichment of individual binders (L10 and L11) from the RFdiffusion 5 campaign across increasing rounds of MACS. **(f)** Biolayer interferometry (BLI) curves and dissociation constants (Kd) values for L10 and L11 binding to BCMA antigen (100nM) from the RFdiffusion 5 campaign. **(g)** Enrichment of individual binders from BindCraft1 (BC1) across successive rounds of magnetic cell sorting (MACS) at decreasing concentrations, demonstrating progressive selection of high-affinity binders. **(h)** BLI curves and estimated Kds for additional sequencing hits from the BC1 campaign.

**Extended Data Figure 2.**
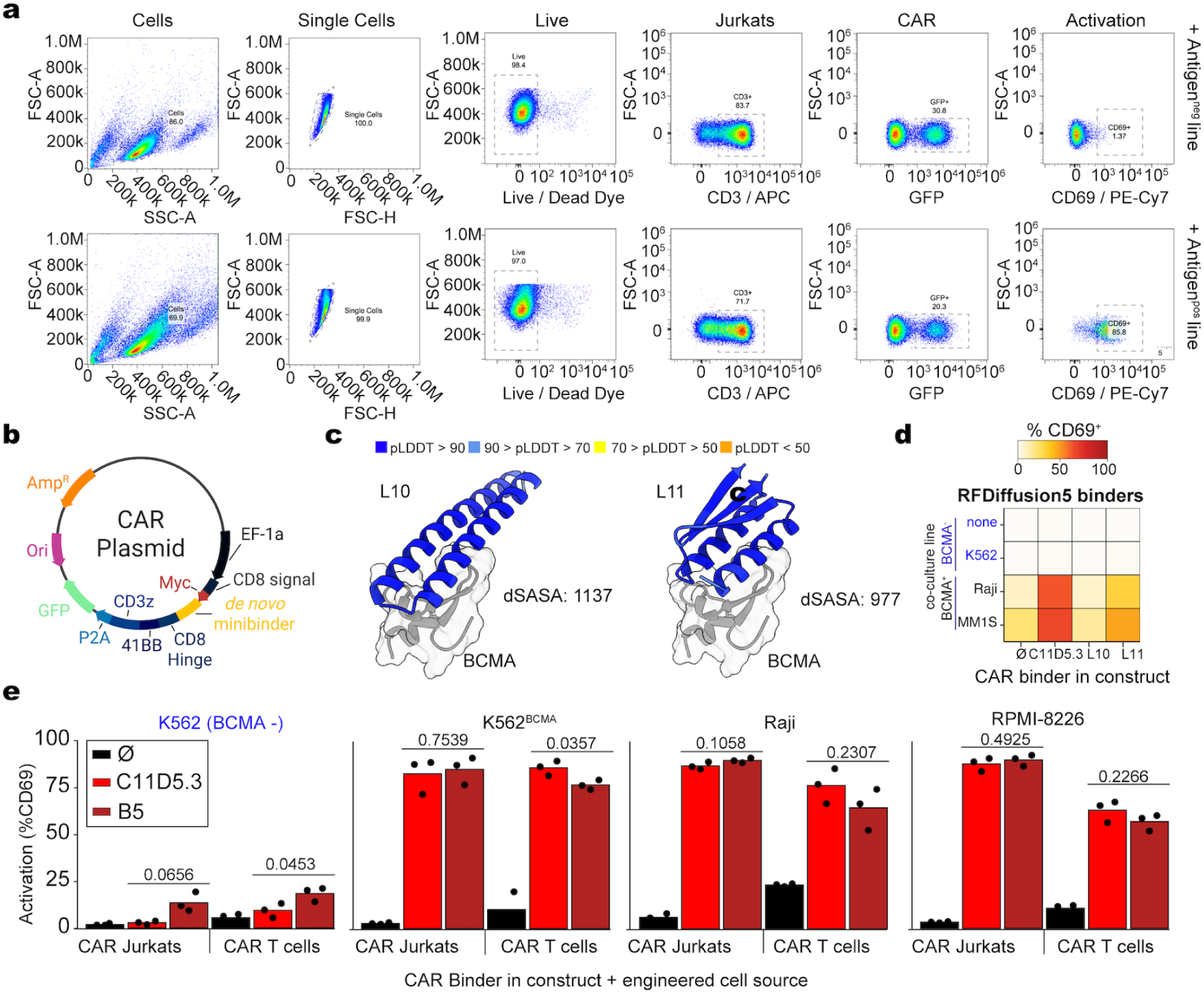
Characterization of lead *de novo* designed BCMA binders. **(a)** Representative gating strategy for CAR Jurkat and CAR T cell co-culture assays used to quantify proportion of cells with CD69 activation. **(b)** CAR construct used in CAR Jurkat and CAR T assays containing an EF1α promoter, CD8 signal peptide, *de novo* minibinder, CD8 hinge, 4-1BB domain, P2A sequence, and GFP reporter. **(c)** Predicted structures of two *de novo* CAR binders designed by RFDiffusion. Binders are colored by per-residue AlphaFold2 predicted Local Distance Difference Test (pLDDT) metrics. **(d)** CAR Jurkat co-cultures expressing L10 and L11 BCMA-specific *de novo* minibinders with BCMA⁻ and BCMA⁺ cancer cell lines. **(e)** %CD69 activation was quantified for CARs expressing no binder (Ø, black), C11D5.3 (red), or B5 minibinder (dark red), in either Jurkat cells or primary T-cells in co-cultures with CD19⁻ and CD19⁺ cancer cell lines, comparing Jurkat and primary CAR T responses within each binder condition. *P* values calculated using Student’s *t*-test.

**Extended Data Figure 3.**
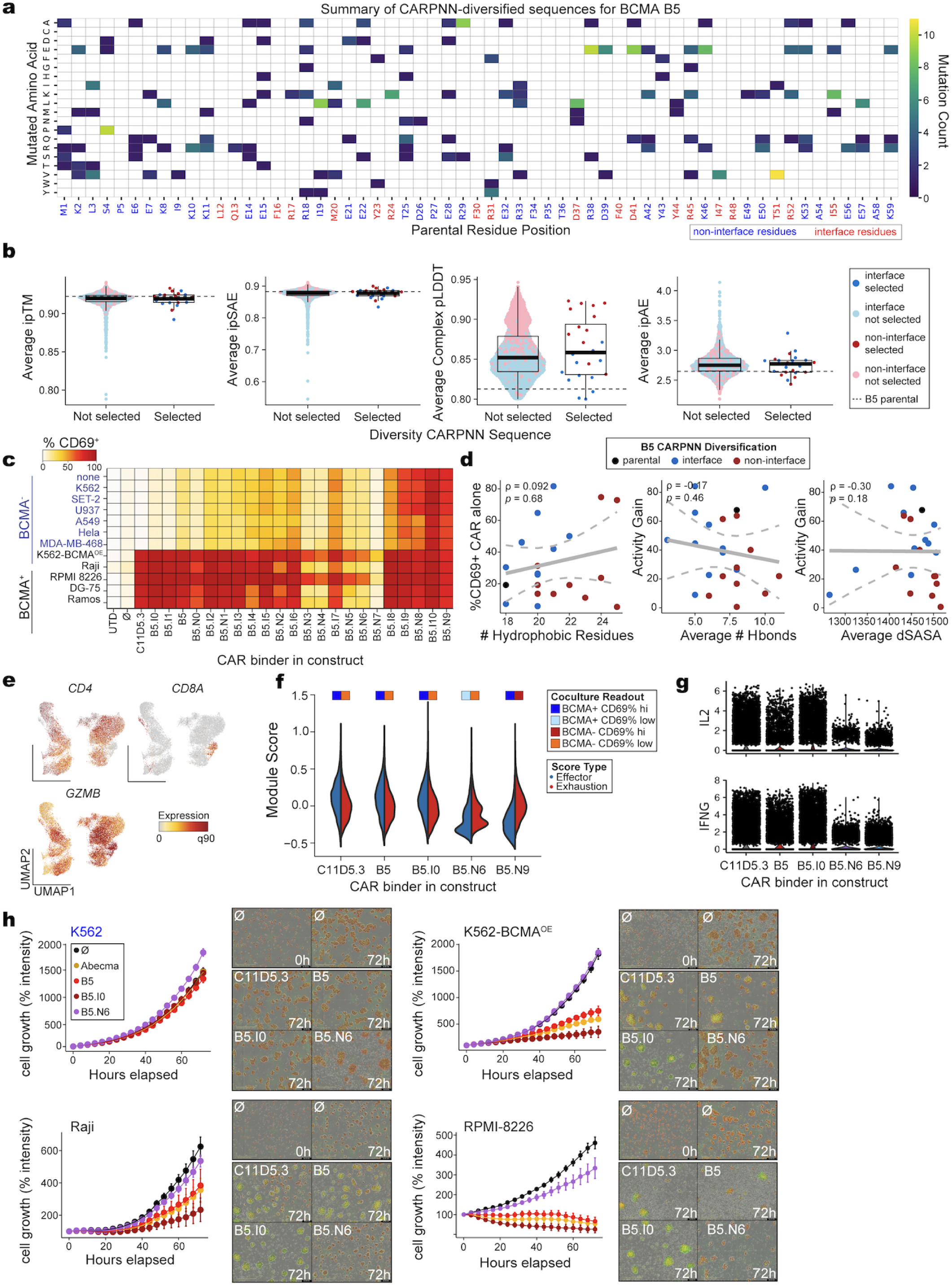
Characterization of the BCMA evolution campaign. **(a)** Summary of mutations introduced to each of the diversified BCMA binders relative to the parental B5 binder. **(b)** Design metrics of candidate and selected binders from sequence diversification of the BCMA binder. The parental B5 value is noted in the dashed line in each panel. **(c)** Summary of diversified sequences as CAR-Jurkats in co-cultures with variable cell lines to assess potential off-target activation and CAR specificity. **(d)** Association between attributes diversified during the CARPNN mutagenesis experiment and the CAR attributes they were hypothesized to influence. **(e)** Marker gene expression across CAR T cells. **(f)** Effector and exhaustion module score across CAR T cells and their relative CD69% in coculture experiment. **(g)** Single-cell expression of key effector genes across variable CAR T binders. **(h)** Incucyte killing assays showing the cytolytic activity of CAR T cells expressing either BCMA-specific minibinder- or scFv-based receptors. Time-course plots showing normalized red calibrated unit (%RCU) intensity relative to time 0h for each construct (left). Representative images at 0h and 72h for empty binder and at 72h for each binder condition to illustrate target-cell killing (right). Green fluorescence denotes CAR T cells, and red fluorescence denotes the corresponding target cell line.

**Extended Data Figure 4.**
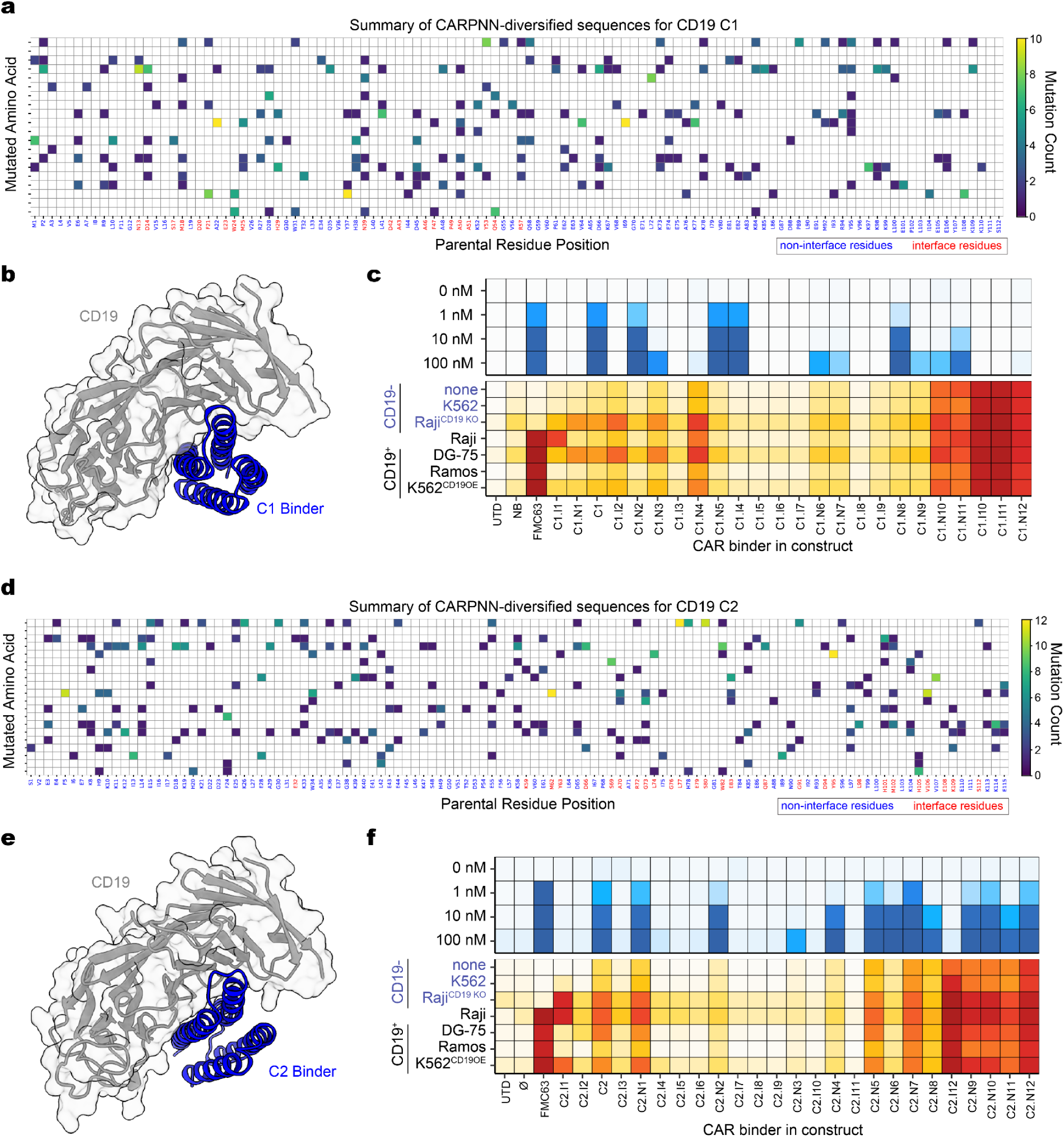
Characterization of the CD19 evolution campaign. **(a)** Summary of mutations introduced to each of the CARPNN diversified CD19 C1 binder. Red residue index denotes interface residues while blue index denotes non-interface residues. **(b)** Predicted structure of the C1 binder in complex with CD19. **(c)** Heatmap of the mutagenized C1 binder variants tested as CARs in co-cultures. Upper panel (blue) displays CAR binding across the indicated antigen concentrations. The lower panel (red) reports %CD69 activation in CD19^-^ and CD19^+^ cell lines. **(d)** Summary of mutations introduced to each of the CARPNN diversified CD19 C2 binder. Red residue index denotes interface residues while blue index denotes non-interface residues. **(e)** Predicted structure of the C2 binder in complex with CD19 **(f)** Same heatmap layout as in **(c)** but shown here for the mutagenized C2 binder variants tested as CARs.

**Extended Data Figure 5.**
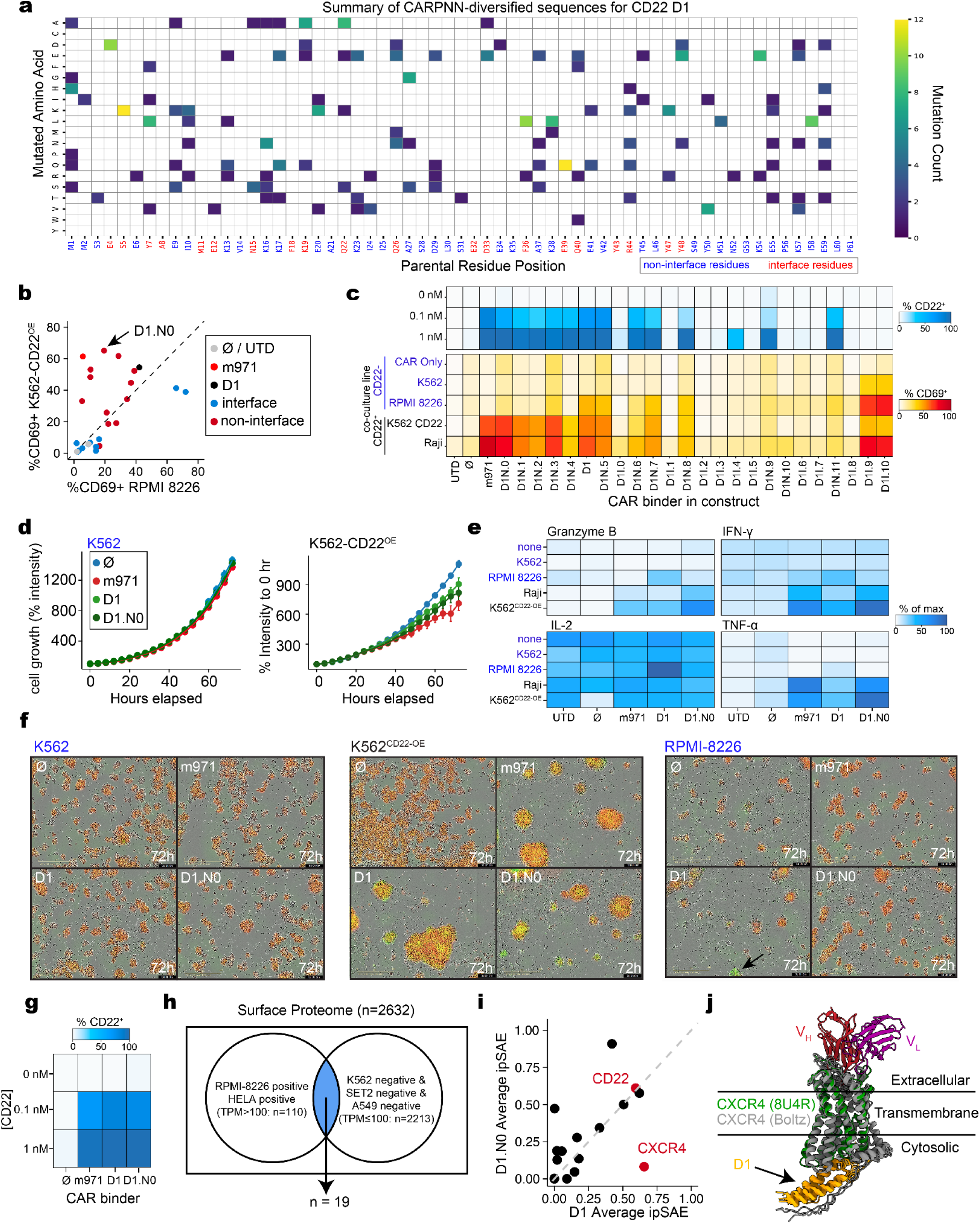
Characterization of the CD22 evolution campaign. **(a)** Summary of mutations introduced to each of the CARPNN diversified CD22 D1 binder. Red residue index denotes interface residues while blue index denotes non-interface residues. **(b)** Comparison of CAR activation of the evolved CD22 D1 binders in CD22^-^ RPMI 8226 cell lines and CD22-overexpressing K562 cell lines. **(c)** Summary of diversified sequences from antigen CAR flow (top) and co-cultures with variable cell lines (bottom). **(d)** Representative Incucyte killing assays showing the cytolytic activity of CAR T cells expressing either CD22-specific minibinder- or scFv-based receptors. Time-course plots showing normalized red calibrated unit (%RCU) intensity relative to time 0h for each construct. **(e)** Cytokine productions from CD22-specific CAR T cells in co-cultures with CD22^+^ and CD22^-^ target cell lines. Heatmap shows mean cytokine levels across triplicates, revealing elevated cytokine release specifically in response to CD22-expressing targets, consistent with antigen-specific activation and killing. **(f)** Representative images at 0h and 72h for NB and at 72h for each binder condition to illustrate target-cell killing. Green fluorescence denotes CAR T cells, and red fluorescence denotes the corresponding target cell line. **(g)** Characterization of CAR antigen binding at variable CD22 concentrations. **(h)** Identification of plausible candidates of D1 off-target interaction via subsetting HPA surfaceome and GTEx overlap. **(i)** Comparison of average cofolding ipSAE score between parental D1 to all plausible off-target genes and the evolved D1.N0 binder to plausible off-target genes. **(j)** Predicted binding site of parental D1 binder towards CXCR4 aligned to a solved structure of CXCR4 (PDB: 8U4R).

**Extended Data Figure 6.**
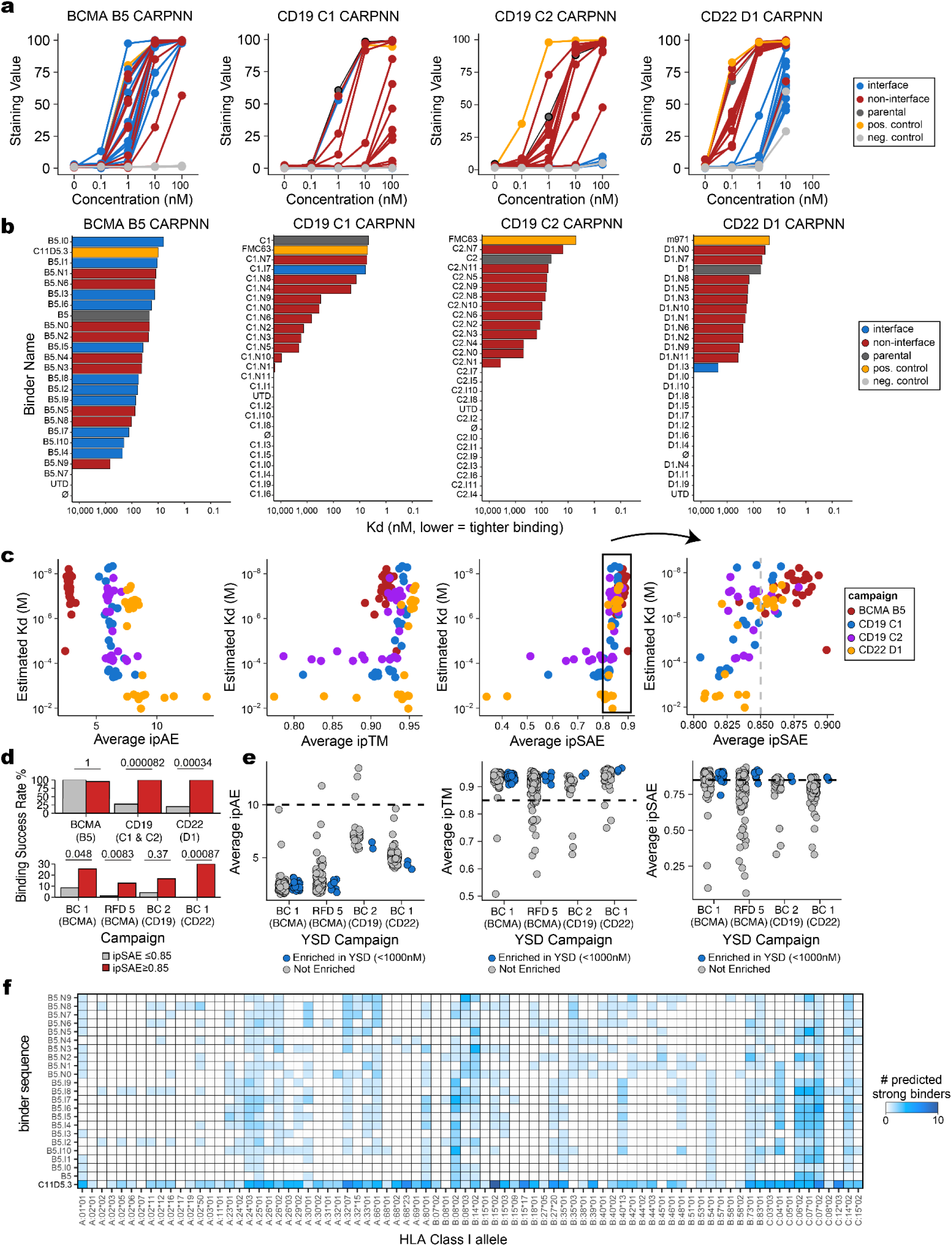
Additional assessment of campaign success rate and binder qualities via *in silico* metrics. **(a)** Staining intensity for CAR Jurkats with antigen at various concentrations. **(b)** Binding affinity estimates of all assessed CARPNN diversified binders based on non-linear least squares (NLS) fit of the staining values for good fits (*R*^2^ > 0.95) or single point estimates. **(c)** Comparison of various protein interaction metrics and their relation to estimated affinity across campaigns. The rightmost panel is a zoomed-in view of the relationship between estimated affinity and average ipSAE. **(d)** Comparison of binding success rate of the CARPNN and YSD campaigns using various interaction metrics as cutoffs. CARPNN campaign binding success is defined as having estimated affinity below 1000nM; YSD campaign binding success is defined as being enriched in the 1000nM sorted population (CPM > 1000, CPM fold change against parental population > 2). **(e)** Comparison of *in silico* metrics across yeast surface display (YSD) campaigns where binders were primarily designed using RFdiffusion (RFD) or BindCraft (BC). A sequence is considered YSD enriched at 1000nM if it has at least 1000 CPM (Counts per Million) at 1000nM and the CPM is at least two-fold higher than its CPM in the unsorted parental pool. **(f)** Comparison of the number of strong HLA-Class I binding peptides of all evolved BCMA binders compared to C11D5.3 (scFv in Abecma) across diverse HLA types.

## METHODS

### RFdiffusion

To generate *de novo* binders against tumor-associated antigens, we first used RFdiffusion to generate protein binder backbones against the crystal structure of the extracellular domain of BCMA bound to its native ligand APRIL (PDB: 1XU2). We launched three campaigns targeting two sets of hotspots consisting of hydrophobic residues: one campaign targeting the outward-facing leucines in positions 17, 26, and 35 (BCMA RFdiffusion Campaign 1), and another two sets of campaigns targeting hydrophobic residues consisting of the phenylalanine at position 14, leucine at position 18, and alanine at position 21 (BCMA RFdiffusion Campaign 2, BCMA RFdiffusion Campaign 3). In all campaigns, we generated protein backbones with lengths ranging between 30-90 without conditioning on any structural scaffolds. Backbones with no alpha-carbon less than 10Å away from BCMA were also discarded.

After failing to yield high-affinity binders in our initial campaigns, we sought to target new sets of epitopes that may be more conducive to protein-protein interaction. Using PeSTo^51^, a deep-learning model that predicts protein-binding surfaces on biomolecules, we identified three residues predicted to have a high likelihood of forming protein-protein interactions on BCMA (1XU2, Chain R), leucine17 (score = 0.95), histidine18 (score = 0.9), and alanine19 (score = 0.88) (BCMA RFdiffusion Campaign 4).

For all RFdiffusion campaigns, we used the filtered backbones as input into ProteinMPNN^21^ with recommended settings^9^ and generated 10 putative sequences assigned per backbone. To identify predicted high-confidence binders, we filtered via the “AF2 initial guess” protocol^9^, which enables fast screening of binders. In brief, the putative binder sequences are refolded against the target structure, and high-quality interactions were filtered such that the resulting co-fold had a complex pLDDT > 80 and pAE interaction < 10. Binders passing *in silico* metrics were then sorted by ascending pAE interaction, and the top 100 sequences with the lowest pAE interaction per backbone were selected for screening via yeast display.

For CD19, we applied the same RFdiffusion-ProteinMPNN-AF2 pipeline to generate 100 protein binders against the crystal structure of the extracellular domain of CD19 bound to FMC63 (PDB: 7URV), with hotspots selected to be isoleucine at position 73, valine at position 241, and leucine at position 247 (CD19 RFdiffusion Campaign 1). Notably, the *in silico* success rate of backbone generation against CD19 was much lower than that of BCMA, likely in part due to the much larger target structures. Next, we used a similar PeSTo-based approach to identify four residues on CD19 (PDB 7URV, chain C) with high probability for protein interaction: leucine188 (score = 0.81), methionine190 (score = 0.85), threonine195 (score = 0.89), and tryptophan197 (score = 0.88) (CD19 RFdiffusion Campaign 2). These residues are then provided as hotspots to RFdiffusion, and the same selection strategy was applied to generate 100 CD19 binders for experimental validation.

After the second round of BCMA and CD19 campaigns failed to yield high-affinity binders, we hypothesized that the lack of binding was not due to insufficient contact with the target but rather the lack of stable binder structures. To test this hypothesis, we removed the backbone diversity filter and decided to constrain the overall fold of the binders to previously validated protein scaffolds^8^ as provided by the RFdiffusion developers and used the fold-conditioning protocol to generate 134 binders against BCMA (BCMA RFdiffusion Campaign 5) and 196 binders against CD19 (CD19 RFdiffusion Campaign 3, CD19 RFdiffusion Campaign 4) with the same epitope specification and selection strategy as previous campaigns. A moderate affinity BCMA binder was generated using this strategy, while no CD19 binder was obtained.

For CD22, a similar strategy was used. Using the crystal structure of the Ig domains 1-3 of CD22 (PDB: 5VKJ, chain A^52^) as input, we did not identify any residues with high PeSTo scores. Thus, we selected three hydrophobic residues that are close in proximity as hotspot inputs: Leucine 161, Leucine 193, and Phenylalanine 206. A fold-conditioned RFdiffusion campaign was launched with the same selection as the previous campaign to generate 102 CD22 binders (CD22 RFdiffusion Campaign 1).

### BindCraft

Motivated by the success of BindCraft in generating binders against EGFR in a public competition^23^, we launched *in silico* campaigns against BCMA, CD19, and CD22 using BindCraft, providing the same antigen crystal structures provided to RFdiffusion as input. We observed high-confidence binder structure against both antigens without pre-specifying hotspot residues and proceeded to generate the rest of the binders using similar settings. In the BCMA BindCraft campaign (BCMA BindCraft Campaign 1), binders of length 60-65 were specified. The default four-stage multimer setting was used to generate hallucination trajectories; the binders are then subject to the default filters; and the campaign concluded after ∼300 filter passing designs were reached. The filter passing BCMA binders are then ranked by ipTM as recommended, and after deduplicating sequences derived from the same protein backbone, the top 125 sequences were selected for experimental testing.

In the first CD19 BindCraft campaign (CD19 BindCraft Campaign 1), the binders were similarly ranged between 50-65 amino acids, and the default hallucination trajectory settings were used. However, we noticed a low initial generation success rate and experimented with other setting configurations, varying parameters such as removing target template sequence during design and re-prediction (the “default_4stage_multimer_flexible” setting), providing binder template during re-prediction (the “default_4stage_multimer_hardtarget” setting), and increasing the helicity weight penalty (the “betasheet_4stage_multimer_hardtarget” setting). The design binders passing default filters using different settings were then aggregated, and the top 100 binders with the highest ipTM are selected.

For the second CD19 BindCraft campaign (CD19 BindCraft Campaign 2), we reasoned that the lack of binding was due to insufficient contact surface between the de novo binder and CD19. As such, we adjusted the generation setting to binders between length 100-200 and used similar filters and diverse trajectory settings as the first CD19 BindCraft campaign. For this campaign, we decided to implement a more stringent pLDDT filter of 0.9 and proceeded with 30 binders with the highest pLDDT. The lower selected binder number was to accommodate synthetic DNA orders as eBlocks rather than o-Pools to minimize costs.

To design binders against CD22, we reasoned that targeting an epitope that is more membrane proximal may increase CAR efficacy^41^. As such, we decided to use the crystal structure of the membrane proximal d6-d7 domain of CD22 bound to the pre-clinical anti-CD22 scFV m971 (PDB: 7O52). Given our success in generating BCMA and CD19 binders using BindCraft, we used similar hallucination trajectory settings and filters to generate ∼500 *in silico* binders between length 50-65 and picked the top 100 binders ranked by ipTM for experimental testing (CD22 BindCraft Campaign 1).

### Lead optimization with CARPNN

To optimize our lead binders while preserving most of their functional properties as CARs, we reasoned that diversifying a limited set of residues while maintaining high confidence *in silico* metrics may help us explore the functional landscape more efficiently. To perform sequence variation on the parental structure, we divide a binder’s residues into interface or non-interface residues based on whether any atoms of the residue are within 4Å of any atoms in the target structure. Each set of the residues is then mutagenized via SolubleMPNN with a temperature parameter of 0.4, with 2000 draws each (BCMA) or 5000 draws each (CD19, CD22). A Levenshtein distance cutoff of 5 (BCMA) or 10 (CD19, CD22) is then applied to ensure the mutated binder sequences are sufficiently different in sequence composition from the parental binder. After the sequences are selected, these sequences are then folded against the target antigen using a BindCraft-based Boltz filtering pipeline. In particular, we folded the candidate binder sequences against the target antigen sequences using Boltz1 with 5 diffusion samples and 10 recycles. Key metrics such as pLDDT, pAE, pLDDT and other biophysical metrics were then calculated and averaged across the 5 models. Filter terms used in BindCraft were then converted to their closest equivalent terms reported by Boltz1. For example, the cutoff used for ‘Average_pLDDT’ BindCraft is used to define the cutoff for ‘Average_complex_plddt’ in Boltz1. To ensure the backbone structures are controlled for between binders, a filter is applied to remove binders whose alpha carbon RMSD is more than 1Å away from the parental backbone.

After filtering for high-quality *in silico* binders, we diversified on 4 binder attributes to maximize observing attributes that correlate with CAR activity. For interface mutagenized binders, we performed k-means clustering on the binders by the number of interface hydrogen bonds formed, as well as dSASA against the target antigen. 12 cluster representatives are then selected for experimental testing. For non-interface mutagenized binders, we employed the same strategy and clustered them by the total number of hydrophobic residues and the total charge of the binder at pH 7.4, and similarly selected the cluster representatives for experimental testing.

### Yeast surface display

Yeast display constructs were generated using the pETcon(-) vector (Addgene, 41522), which contains a GAL1 promoter for inducible minibinder expression and TRP1 as the auxotrophic selection marker. The pETcon(-) backbone was linearized using the restriction enzymes NdeI and BamHI. Each design campaign was synthesized as an oligonucleotide pool (oPool, Integrated DNA Technologies) containing 5′ and 3′ homology arms to the pETcon(–) backbone, which had been linearized with NdeI and BamHI.

For each oPool campaign, 100 ng of DNA was PCR-amplified and purified using the QIAquick PCR Cleanup Kit (Qiagen, Cat. 28104). The amplified oPool and the linearized backbone were combined and concentrated using the DNA Pellet Paint (Millipore, Cat. 70748-3). *S. cerevisiae* strain EBY100 was transformed with 4,000 ng of PCR-amplified insert containing pETcon(–) backbone homologous arms and 1,000 ng of linearized vector using the Frozen-EZ Yeast Transformation II™ Kit (Zymo Research, Cat. T2001). Successfully transformed clones were selected using SDCAA growth medium for 48h at 30°C.

For minibinder library display, yeast cells were transferred into SGCAA medium at a density of 0.3–0.5 × 10⁷ cells mL⁻¹ and incubated for 16 h at 20 °C with shaking at 200 rpm. Cells were collected by centrifugation at 4,000 × g for 1 min and washed once with PBS-B buffer (PBS supplemented with 0.1% (w/v) BSA). For binding assays, induced yeast cells were incubated with biotinylated antigen for 1 h at room temperature, washed twice with PBSF, and stained with a cocktail of APC–streptavidin and anti-Myc–FITC antibodies.

Yeast libraries displaying minibinders were induced for 24 h at 20 °C in SGCAA medium with shaking at 200 rpm. Following induction, cells were washed twice with PBS-B (PBS supplemented with 0.1% BSA) and incubated with Biotin-Binder Dynabeads (Thermo Fisher Scientific, Cat. 11047) without antigen to deplete clones exhibiting nonspecific bead binding. The supernatant containing unbound yeast was collected and incubated with Dynabeads pre-conjugated to biotinylated target antigen for positive selection. After incubation, bead–yeast complexes were washed five times with PBS-B to remove unbound cells and resuspended in 50 mL of SDCAA medium for outgrowth and enrichment of antigen-binding clones.

### Preparation of yeast libraries for sequencing

Plasmid DNA was isolated from yeast using a modified QIAGEN Plasmid Midi Kit (Qiagen, Cat. 12143) protocol. Briefly, 2g of yeast cells were harvested and converted to spheroplasts in SCE buffer (1M sorbitol, 0.1 M sodium acetate, 60mM EDTA, pH 7.0) with zymolyase (5 mg) and β-mercaptoethanol (30 µL) for 1 h at 37 °C. Spheroplasts were then lysed, purified, and the plasmid was eluted following the manufacturer’s protocol. Enriched plasmid libraries were modified by PCR amplification to append sequencing-compatible adapter elements with a forward primer pool 5’-TCGTCGGCAGCGTCAGATGTGTATAAGAGACAG(N)_0-2_GGTCGGCTAGCCATATG-3’ and reverse primer pool 5’-GTCTCGTGGGCTCGGAGATGTGTATAAGAGACAG(N)_0-2_CTTTTGTTCGGATCC-3’. PCR amplicons were purified using SPRI bead cleanup, and Illumina i5 and i7 index sequences were subsequently appended to generate sequencing-ready libraries. Final libraries were quantified and size-verified by TapeStation (Agilent) prior to dilution and loading onto the Element Aviti sequencer. Library abundance was determined by building a custom index of DNA sequences using kallisto^53^, and each library was quantified using default parameters for a paired-end sequencing library.

### Recombinant binder expression and purification

Minibinder sequences were cloned into a modified pBAD bacterial expression vector using Gibson assembly, incorporating a C-terminal 6×His tag. Each construct was transformed into chemically competent *E. coli* BL21 (DE3) cells (Invitrogen, Cat. C600003) and plated on LB-agar containing 100 µg mL⁻¹ carbenicillin. Single colonies were inoculated into 50 mL of Terrific Broth (TB) and cultured overnight at 37 °C. The following day, 2 mL of the starter culture was transferred into 1 L of TB and grown at 37 °C until reaching an optical density at 600 nm (OD₆₀₀) of 0.6–0.8. 20% arabinose (Fisher Scientific, Cat. AAA1192130) was then added to a final concentration of 0.2% (w/v) to induce minibinder expression, and cultures were incubated overnight at 37°C with shaking. The following day, cells were collected by centrifugation at 4,000 RPM for 10 minutes, resuspended in lysis buffer supplemented with protease inhibitor tablets (Pierce Cat. A32963) (50mM Tris-HCl (pH 8.0), 300mM NaCl, 15mM imidazole, 0.25% NP-40, 0.25% Triton, and 10% glycerol), and sonicated for 1 minute at 1 sonication per 3 seconds (3x with a 3 minute break in between sonication). Whole-cell lysates were clarified by centrifugation at 18,000 rpm for 1 h at 4 °C. Minibinder proteins were purified from the supernatant using His60 Ni Superflow Resin (Thermo Fisher Scientific, Cat. A50589) and washed with buffer (50 mM Tris-HCl, 300 mM NaCl, 40 mM imidazole, 10% glycerol), followed by elution with buffer (50 mM Tris-HCl, 300 mM NaCl, 250 mM imidazole, 10% glycerol) using a gravity-flow column. Eluted proteins were purified by size-exclusion chromatography on a Superdex 75 Increase 10/300 GL column (Cytiva, Cat. 29148721) in PBS (pH 8.0).

### Biolayer Interferometry

Binding kinetics were measured using the Gator Pro Biolayer Interferometry (BLI) system (Gator Bio). Streptavidin biosensors (GatorBio, Cat. 160002) were equilibrated in PBS-B (PBS supplemented with 0.2% BSA) and loaded with biotinylated target antigen (20 nM) during the loading phase. Following a brief baseline step, biosensors were transferred into wells containing minibinder proteins at increasing concentrations to monitor association (400s), then into buffer-only wells to measure dissociation (600s). Sensorgrams were reference-subtracted and fitted using the Gator Pro analysis software to derive kinetic parameters (k_on_, k_off_) and equilibrium dissociation constant (K_d_).

### CAR Jurkat cell screening

The CAR lentiviral backbone (PWL003) was constructed with a CD8a signal peptide and hinge region, followed by a *de novo* designed minibinder sequence flanked at the N terminus by a Myc-epitope tag for detection driven by an EF1α promoter within a second-generation lentiviral cassette. The intracellular domains consisted of a 4-1BB and CD3ζ to provide costimulatory and activation signaling, respectively.

For engineering Jurkats, lentiviral particles were generated by co-transfecting HEK293T cells with the PWL003 lentiviral vector, the packaging plasmid psPAX2, and the envelope plasmid VSVG using FuGENE HD transfection reagent (Promega, Cat. E2311). Specifically, 6×10^6^ HEK293T cells were seeded in 10 cm dishes containing 10 mL DMEM supplemented with 10% FBS and 1% penicillin–streptomycin. On the same day, 6 µg of PWL003, 3 µg of psPAX2, and 1 µg of VSVG were combined and mixed with 500 µL Opti-MEM and 18 µL FuGENE HD. After 15 minutes of incubation at room temperature, the mixture was added dropwise to the HEK293T cells. At 18 hours post-transfection, the medium was replaced with 10 mL of fresh DMEM. Lentiviral supernatants were collected 48 hours after transfection and filtered through a 0.45 µm syringe filter (09-754-21, Fisher Scientific).

For transduction, 5 × 10⁵ Jurkat cells were seeded in 2 mL of RPMI medium per well of a 6-well plate, and three viral titers (100–800 µL of freshly harvested lentiviral supernatant) were applied. Polybrene (4 µg mL⁻¹) was added, followed by spin inoculation at 800 × g for 30 min. Transduction efficiency was assessed 60 h post-infection by flow cytometry as the percentage of GFP^+^ cells. CAR Jurkat populations exhibiting 25–55% GFP positivity were selected for subsequent experiments.

### CAR Jurkat co-culture CD69 activation assays

CAR Jurkat (CAR J) cells were selected based on GFP positivity, counted, and centrifuged at 300 × g for 5 minutes. Cells were resuspended in RPMI medium at 5 ×10^5^ live GFP^+^ cells/mL. Since GFP positivity varied across transductions, the total number of cells used for resuspension was adjusted accordingly to achieve this target concentration of GFP^+^ cells. Target cell lines, either expressing or lacking the cognate surface antigen, were also counted and adjusted to 5 ×10^5^ cells/ mL in RPMI medium. Co-cultures were established in 96-well u-bottom plates by seeding 5 ×10^4^ GFP^+^ CAR J cells with 5 ×10^4^ target cells per well. For co-cultures with variable effector : target (E:T) ratio, 2.5 ×10^4^ GFP^+^ CAR J cells were seeded in each well, and the number of target cells is adjusted accordingly. Control wells containing CAR J cells alone or target cells alone were included. Co-cultures were incubated for 18 hours at 37 °C in the incubator.

After the incubation, the co-culture plate was centrifuged at 300 × g for 5 minutes. The supernatant was discarded, and the cells were washed with 200 µL of PBS containing 0.5% BSA. The cells in each well were then stained with 100 µL of the antibody cocktail diluted in PBS containing 0.5% BSA consisting of 0.1 µL of LIVE/DEAD Fixable Near IR (780) stain (Invitrogen, Cat. L34992), 0.5 µL of anti-CD3 APC conjugated antibody,(Biolegend, Cat. 300312), 0.5 µL of anti-CD69 PE-Cy7 conjugated antibody (Biolegend, Cat. 310912), and 0.5 µL of anti-target antigen fluorophore conjugated antibody (**Extended Data Table 2**). After 30 minutes of incubation at 4°C, the cells were washed once and resuspended in 150 µL of PBS with 0.5% BSA. The 96-well plate containing the cell samples was then processed by Attune CytPix Flow Cytometer and CytKick Autosampler (Invitrogen, Cat. A42901).

### Recombinant antigen binding assay for CAR-Jurkats

CAR Jurkat cells (25,000 per well) were seeded in 96-well plates in 100 µL of RPMI-1640 medium supplemented with 10% fetal bovine serum (FBS) and incubated overnight (16 hours) at 37 °C. The following day, cells were washed once with 1× PBS-B (0.25% BSA) and incubated with varying concentrations of biotinylated recombinant antigen BCMA (Sino Biological, Cat. 10620-H40H-B), CD22 (Sino Biological, Cat. 11958-H49H-B), or CD19 (Sino Biological, Cat. 11880-H49H-B) for 1 hour at room temperature. After incubation, cells were washed twice with 1× PBS-B and stained with anti-Streptavidin-APC conjugated antibody (BioLegend, Cat. 300312) for 30 min at 4°C. Cells were then washed twice and resuspended in 150 µL of PBS-B for flow cytometry analysis.

### Primary CAR T assays

T cells were isolated from Stem Cell Leukopacks (Stem Cell, Cat. 200-0092) using EasySep Human T Cell Isolation Kit (Stem Cell, Cat. 17951). Upon thawing, cells (Day 0) were resuspended in Immunocult (Stem Cell, Cat. 10981) supplemented with 0.1ug/mL of IL-2 (Stem Cell, Cat. 78036) and activated with ImmunoCult Human CD3/CD28 T Cell Activator (Stem Cell, Cat. 10971). Cells were maintained at a concentration of 0.25e6/mL. On Day 2, concentrated lentivirus was added to cells, and cells were spin-infected at 800g for 45 minutes. On Day 5, DAPI-stained T cells were sorted for GFP^+^ cells using the SH800S Cell Sorter.

On Day 11, cells were collected to set up co-cultures, for flow: 50,000 each of target and effector cells were resuspended in 200ul per well of a 96-well plate, incubated for 16 hours, for incucyte:10,000 each of target and effector cells were resuspended in 200ul per well of a 96 well plate. Cells in the flow co-culture plate were spun down, and supernatant was collected and frozen at -80°C in 10 μL aliquots for ELISA assays, and cells were processed for analysis via flow cytometry.

For ELISA, the supernatant was prepared according to the manufacturer’s instructions. Kits and input of supernatant used: IFN-y (Biolegend, Cat. 430101), IL-2 (Biolegend, Cat. 431801), Granzyme B (Invitrogen, Cat. BMS2027-2), TNF-α (Biolegend, Cat. 430201).

For live-cell cytotoxicity assays, cells were co-seeded in 200 µL of medium per well in an IncuCyte ImageLock 96-well microplate (Sartorius, Cat. BA-04856). Target cells were stably labeled with mCherry, and CAR-T cells were labeled with GFP. Live-cell imaging was performed using the IncuCyte system, acquiring four fields per well every 4 h for 72 h. Cell viability was quantified as the total red object integrated intensity (RCU × µm² per image) normalized to the initial time point (0 h).

### Single-cell RNA-seq

CAR T cells were co-cultured with target antigen–positive or target antigen-negative cells for 24 h, harvested, and enriched by FACS for CD3⁺ T cells. Cells were washed in Cell Staining Buffer (Biolegend, Cat. 420201), blocked with TruStain FcX (BioLegend, Cat. 422301) for 10 min at 4 °C, and stained with hashtag oligonucleotide (HTO) antibodies (0.25 µg per sample; TotalSeq-C, as appropriate) for 30 min at 4 °C. After two washes, individual CAR T populations were counted, assessed for viability, and combined at the desired ratios for a minimum of 30k total cells.

Single cells were partitioned into GEMs (gel bead-in-emulsions) on a Chromium Controller (10x Genomics; Single Cell 5′ kit, Cat. PN-1000695) following the manufacturer’s protocol with Feature Barcoding for hash tag oligos (HTOs; 10x Genomics, Cat. 1000703). Reverse transcription was performed in-GEM, followed by emulsion breakage, cDNA cleanup, and amplification. Separate gene-expression and HTO libraries were constructed per 10x recommendations. Library quality and size distributions were verified by TapeStation (Agilent) and concentration by Qubit (Thermo). Pooled libraries were sequenced on an Element AVITI instrument using paired-end chemistry with 10x Genomics compatible read lengths. Different CARs were multiplexed with 5’-compatible Totalseq-C reagents. Per-cell hash quantifications were estimated using kallisto|bustools^54^ for the relevant hashing antibodies, and per-cell hash assignments were annotated using default Seurat settings^55^.

After demultiplexing and hashing assignment, each sample’s raw count matrix was loaded as Seurat objects^56^ for downstream analysis. Filtering based on total RNA count, unique RNA detected, and proportion of mitochondria reads was performed to remove low-quality cells. For each sample, read counts were normalized and scaled, and dimensionality reduction was performed to identify cell neighborhoods, followed by Leiden clustering. Cells in clusters lacking *CD3E* expression were considered contaminants and were removed, after which the filtered sets of cells were identically reprocessed in Seurat. To account for cell cycle effects, cell cycle scoring was performed, and Harmony^57^ was applied to produce a phase-regressed embedding. Differentially expressed genes were identified using a default implementation of the Wilcoxon rank sum test between different binders with p-values adjusted for multiple hypothesis testing via the Benjamini-Hochberg procedure. Genes overexpressed by cells expressing the evolved BCMA binder B5.I0 compared to cells expressing the parental BCMA binder B5 were then filtered using a log2 fold-change threshold above 0.25 and adjusted p-value below 0.05 to obtain the minibinder activation gene set. The activation module scores for CD22 experiments were then calculated using this gene set via the AddModuleScore function from Seurat.

### CD22 *in silico* off-target screen

To identify the off-target interactions we observed from the initial CD22 D1 binder, we reasoned the off-target interaction may be mediated by the binder binding to surface proteins that are sufficiently expressed in CD22^-^ cell lines where the CAR is activated but are not sufficiently expressed in CD22^-^ cell lines where the CAR is not activated. Using the curated membrane protein list from the Human Protein Atlas (*n*=2,632) and the cell lines expressions from DepMap, we identified 19 candidates with that are positive for RPMI-8226 and HeLa but are negative for K562, SET2, A549 using either 100 or 1,000 TPM as the cutoff. These 19 candidate off-target proteins were then folded against D1 and the rescued binder D1.N0 via Boltz. Among all predicted interactions, only CXCR4 had significantly higher ipSAE in D1 compared to the evolved D1.N0. However, upon inspection of the predicted structure, in all 5 diffusion samples of the predicted D1-CXCR4 complex the interaction is occurring through the cytosolic end of CXCR4 that is inaccessible to the cell surface (PDB: 8U4R), which suggest the predicted interaction is likely spurious.

### Retrospective analyses of CAR campaigns

To infer the on-cell binding affinities of various binders toward their respective target antigen, we performed non linear least square (NLS) curve fit of the staining intensities at 0.1nM, 1nM, 10nM, 100nM concentrations of the target antigen. The estimated affinities are then rescaled by the ratio between the estimated affinity of the positive control to the reported affinity of the positive controls reported in literature. For curves that have poor fit (R^2^<0.95), a single point estimate based on the staining intensity at the concentration with the highest dynamic range (BCMA: 1nm, CD19 C1: 100nM, CD19 C2: 1nM, CD22 0.1nM) is applied instead and rescaled.

To identify the binder design success rate by yeast binding in an unbiased manner, we defined a binder to be YSD enriched at a specific concentration (*X*) if its CPM from deep sequencing exceeded at least 1000 at the *X* nM sorted pool and it is at least a two-fold enrichment compared to its CPM in the pre-sorted parental pool.

To calculate the predicted class I epitope burden, we computed the predicted affinity between all possible 9-mers of the *de novo* proteins against all supported alleles by NetMHC^44^. The NetMHC Class-I Burden is then computed as the mean fraction of 9-mer peptides predicted to be strong binders for MHC Class I alleles across all alleles supported by NetMHC4.2.

### Statistical analyses and plotting

All data figures were created using the R software environment (version 4.4.1) and ggplot2 (v3.5.1).

